# The Roles of APOBEC-mediated RNA Editing in SARS-CoV-2 Mutations, Replication and Fitness

**DOI:** 10.1101/2021.12.18.473309

**Authors:** Kyumin Kim, Peter Calabrese, Shanshan Wang, Chao Qin, Youliang Rao, Pinghui Feng, Xiaojiang S. Chen

## Abstract

During COVID-19 pandemic, mutations of SARS-CoV-2 produce new strains that can be more infectious or evade vaccines. Viral RNA mutations can arise from misincorporation by RNA-polymerases and modification by host factors. Analysis of SARS-CoV-2 sequence from patients showed a strong bias toward C-to-U mutation, suggesting a potential mutational role by host APOBEC cytosine deaminases that possess broad anti-viral activity. We report the first experimental evidence demonstrating that APOBEC3A, APOBEC1, and APOBEC3G can edit on specific sites of SARS-CoV-2 RNA to produce C-to-U mutations. However, SARS-CoV-2 replication and viral progeny production in Caco-2 cells are not inhibited by the expression of these APOBECs. Instead, expression of wild-type APOBEC3 greatly promotes viral replication/propagation, suggesting that SARS-CoV-2 utilizes the APOBEC-mediated mutations for fitness and evolution. Unlike the random mutations, this study suggests the predictability of all possible viral genome mutations by these APOBECs based on the UC/AC motifs and the viral genomic RNA structure.

**One-sentence summary:** Efficient Editing of SARS-CoV-2 genomic RNA by Host APOBEC deaminases and Its Potential Impacts on the Viral Replication and Emergence of New Strains in COVID-19 Pandemic

## INTRODUCTION

The causative agent of the COVID-19 pandemic, the severe acute respiratory syndrome coronavirus-2 (SARS-CoV-2), is a member of the enveloped Coronaviridae family that has a single-stranded positive-sense RNA genome (*1, 2*). Unlike most RNA viruses that exhibit high mutation rates ((*3*) and references therein), SARS-CoV-2 and other coronaviruses have moderate genetic variability because they have built-in proofreading mechanism in their RNA-dependent RNA-polymerase (RdRP) to correct the errors during viral genome replication and transcription (*4*).

However, sequencing data of SARS-CoV-2 from patients revealed persistent accumulation of new mutations over time, leading to the emergence of new viral strains that can be more transmissible or virulent (*5, 6*) or evade vaccines (*7, 8*), highlighting the importance of understanding the driving force of viral mutation and evolution of SARS-CoV-2 genome. There are two possible main sources for SARS-CoV-2 viral mutations: spontaneous random errors that are not corrected by the build-in proofreading mechanism of the RdRP and the host-factor mediated mutations of viral genome ((*3*) and references therein). Recent analysis of SARS-CoV-2 and rubella vaccine virus genome data derived from patients show predominant mutational patterns with specific signatures rather than random genetic variations, suggesting that the host-factor induced mutations play an important role in shaping the viral genomic RNA mutational outcome and evolution (*9-12*).

The host responses that can cause mutations on SARS-CoV-2 include <u>r</u>eactive <u>o</u>xygen <u>s</u>pecies (ROS) ((*13*) and references therein) and two families of human RNA deaminases: ADARs (the <u>a</u>denosine <u>d</u>eaminases <u>a</u>cting on <u>R</u>NA) and APOBECs (the <u>apo</u>lipoprotein-<u>B</u> (*ApoB*) mRNA <u>e</u>diting enzyme, <u>c</u>atalytic polypeptide-like proteins) (*9, 10, 12*). The ROS could oxidize nucleic acids to cause viral mutations, which is proposed to be related to the G-to-U and C-to-A mutations (*3, 14*). The ADAR enzymes modify adenosine to inosine to cause A-to-G mutations in double-stranded RNA (dsRNA), which play important roles in immune regulation ((*15, 16*) and references therein).

The APOBEC proteins are a family of cytosine deaminases that can deaminate cytosine to uracil (C-to-U) in single-stranded nucleic acids and function in a variety of biological processes, including innate and adaptive immune responses to viral pathogens ((*17–20*) and references therein). The seven APOBEC3 subfamilies (including A3A to A3H) are reported to restrict DNA and RNA viruses (reviewed in (*19, 20*)). While most APOBECs use single-stranded DNA (ssDNA) as substrate for C-deamination (reviewed in (*17*)), three APOBECs, APOBEC1 (A1) (*21, 22*), APOBEC3A (A3A) (*23*), and APOBEC3G (A3G) (*24*), are also shown to deaminate certain cellular single-stranded RNA (ssRNA) targets to cause C-to-U editing. Interestingly, the database analysis of SARS-CoV-2 genomic variations showed an overwhelmingly high C-to-U mutation rate, account for about 40% of all single nucleotide variations, which was interpreted as a result of RNA editing by host APOBECs rather than random mutations (*9, 10, 12, 25*). However, there is no report on the direct experimental evidence to demonstrate if APOBECs can edit the SARS-CoV-2 genome and, if so, what is the extend of editing on the viral genome by different APOBECs, and the potential effect of the APOBEC-editing on the virus. In this study, we investigated whether APOBEC proteins can directly edit the RNA sequence of SARS-CoV-2 to generate C-to-U mutations and how such mutations may impact viral replication and viral progeny production in an experimental system.

## RESULTS

### Experimental design for testing APOBEC-mediated editing of SARS-CoV-2 RNA

Our first goal is to design an experimental system to that can efficiently address whether a particular APOBEC enzyme can actually edit SARS-CoV-2 RNA. We adapted our previously reported cell-based RNA editing system (*26*) to examine the ability of APOBEC proteins to edit SARS-CoV-2 genomic RNA to cause C-to-U mutations. Because A1+A1CF, A3A, and A3G are the three APOBEC proteins shown to possess RNA editing activities (*22–24*), we tested each of these three APOBEC proteins for their ability to edit SARS-CoV-2 RNA. Due to technical and budget limitations for the so called “error-free” safe sequencing system (SSS) (*27, 28*), we selected seven 200 nt-long RNA segments across the SARS-CoV-2 genome for the APOBEC-editing study **(Fig. 1A)**. These seven segments are distributed from the 5’ to 3’ end of the genome to include various viral genes, including the 5’-untranslated region (5’UTR).The selected 200 nt viral RNA segments were constructed into DNA reporter vector fused to the C-terminus of eGFP coding sequence that can be transcribed into RNA under a constitutive promoter (**Fig. 1B**). An AAV intron is inserted in the middle of eGFP that can be useful to differentiate the mature mRNA transcript (with the intron spliced out) from the coding DNA containing the intron. A primer annealing to the exon-exon junction on the mature mRNA **(JUNC, Fig. 1B)** can specifically amplify the RNA, but not the coding DNA, by PCR, making it possible to rule out the C-to-U deamination on DNA from the direct C-to-U RNA editing by APOBECs. The reporter vector was co-transfected with the APOBEC editor vector to express the selected APOBEC protein in HEK293T cells **(Fig. 1C)**. Total RNAs were extracted from the cells for cDNA preparation for sequencing using the SSS approach as described below.

**Fig. 1.**
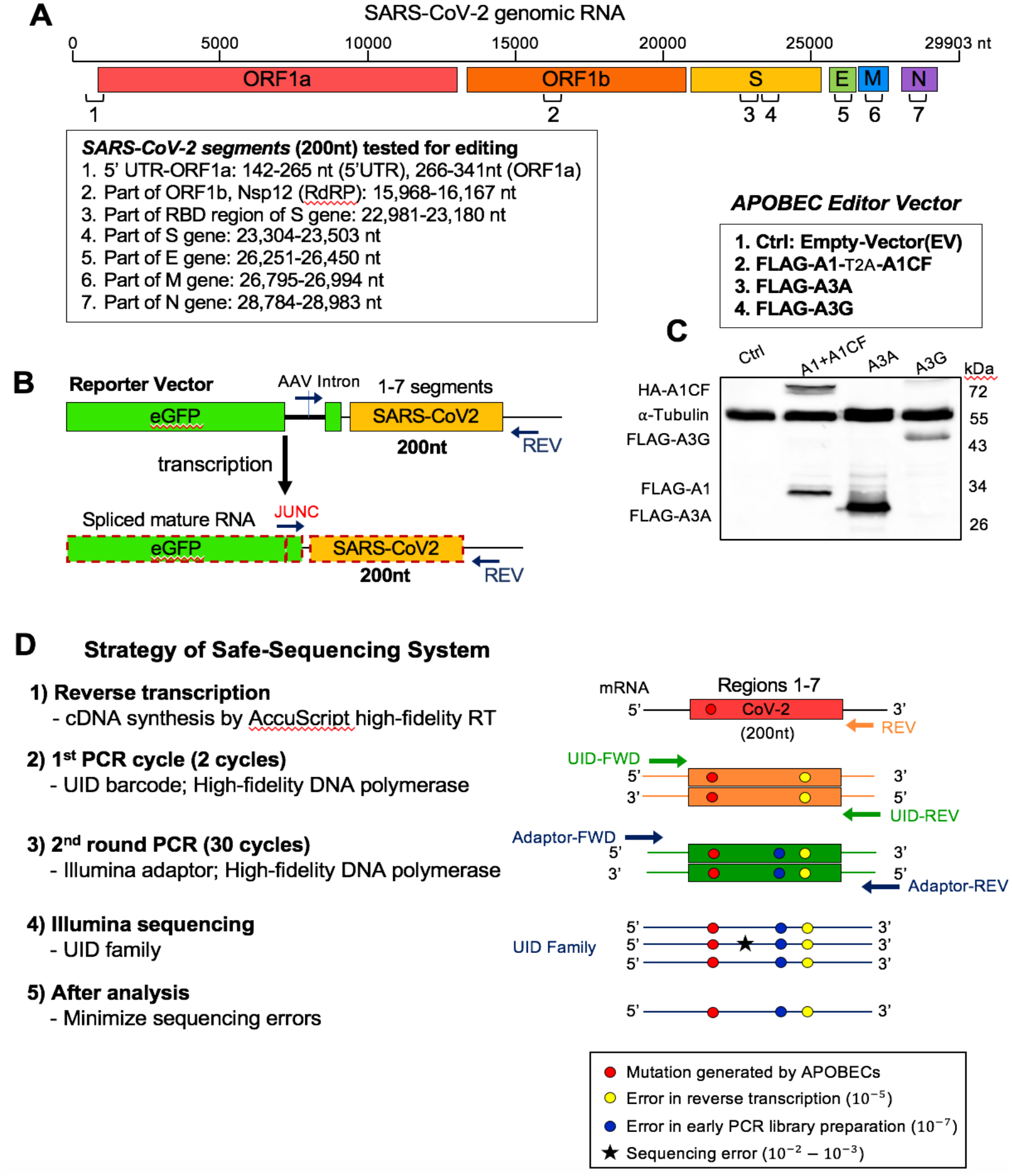
Experimental design of APOBEC-mediated editing of SARS-CoV-2 RNA. **(A)** Diagram of the SARS-CoV-2 genomic RNA, showing the positions (box) of the seven RNA segments (1-7) selected for studying the RNA editing by APOBECs. **(B)** Reporter vector (top) that contain each of the seven selected viral RNA segments that are transcribed into an RNA containing an AAV intron between the eGFP and the viral RNA segment. Splicing out the AAV intron yields a mature mRNA with a new spliced junction sequence (JUNC) that differs from its coding DNA, which can be used to selectively amplify either the mature mRNA or the coding DNA. **(C)** Three APOBEC editor vectors (top, A1-2A-A1CF, A3A, and A3G) and the Western blot showing their expression in 293T cells (bottom). A1-2A-A1CF is constructed as one open reading frame (ORF) with a self-cleavage peptide T2A inserted between A1 and A1CF, which produced individual A1 and A1CF proteins in a 1:1 ratio (*26, 54*). **(D)** Strategy of the Safe-Sequencing-System (SSS) to minimize errors from PCR amplification and sequencing. After the SARS-CoV-2 RNAs from cell extracts are reverse transcribed, the cDNAs are sequentially amplified by the UID barcode (2 cycles) and the Illumina adapter (30 cycles). This SSS approach will distinguish the C-to-U mutations caused by APOBECs from the PCR and sequencing errors (See Methods in SI).

To minimize the sequencing errors when evaluating the C-to-U RNA editing, we employed the SSS system, a targeted next generation deep-sequencing system, with slightly adapted protocols **(Fig. 1D**, see Methods in SI) (*27, 28*). This SSS method involves the following four critical steps. First, the AccuScript high-fidelity reverse transcriptase (known to have ~10^−4^ - 10^−5^ error rates) was used for the initial reverse transcription of the target SARS-CoV-2 RNA transcripts from the cells to single-stranded cDNA. The JUNC forward primer is used to ensure only mature spliced mRNA segments of SARS-CoV-2 were amplified **(Fig. 1B)**. Second, 2 cycles of initial PCR amplification of the cDNA were performed with a both forward and reverse primers containing a Unique IDentifier (UID), a string of 15 nt randomized sequences, to attach the large family of different UID barcodes (~4^30^), discerning each original target molecule. Third, the initial 2 cycle-PCR products were purified and then amplified with Illumina adaptors (PCR error rate is ~10^−7^). Finally, the high rates of errors from paired-end Illumina sequencing (PE150, ~10^−2^ - 10^−3^ error rates) are minimized by eliminating random mutations in the same UID family (see Methods in SI).

Using this SSS approach in this system to check the APOBEC-mediated mutations overcomes two major limitations from directly analyzing the patient-derived SARS-CoV-2 sequences. The first limitation is that the deposited SARS-CoV-2 sequence is “one-consensus-sequence from one-patient”, i.e. only one consensus viral sequence from the NGS sequencing data is reported from one patient sample, which neglects many of the non-consensus sequence variants that may be caused by APOBEC deamination. The second limitation is that regular NGS sequencing’s intrinsic high error rate (PE150, 10^−2^ - 10^−3^ error rates) may drown out the signals of true mutations caused by APOBEC deamination. However, the SSS approach has its own drawback: the SSS sequencing costs several times more than a regular NGS sequencing. Each SSS sequencing read covers ~150-200 nt, making it too expensive for us to cover the entire 30,000 nt RNA genome of SARS-CoV-2. That is the main reason that we selected only seven viral RNA segments for our editing study.

### Sequence motifs near the APOBEC-edited C on SARS-CoV-2 RNA

In our SSS system to identify the APOBEC-edited C-to-U mutations, the average number of UID families for each of the 28 experiments (seven segments each with three different APOBEC enzymes and one control) has an average of about 130,000 (minimum 85,000, maximum 187,000) from a total of ~ 484 million (paired) reads **(Table S1)**. The C-to-U editing levels by each APOBEC are normalized by the control group **(Table S2)**. The C-to-U editing by all three APOBECs is detected, with A1+A1CF and A3A showing much higher editing than A3G **(Fig S1)**. Here, we define the significant target C site where the C-to-U editing efficiency is at least 3 times higher than control. Out of 307 total C in the selected viral RNA segments, the number of significant target sites with A1+A1CF is 135, A3A is 67, and A3G is 11 **(Fig. 2A and Table S2)**. Analysis of the sequence contexts around the significant target C sites (with ± 5 nucleotides from target C) showed that A3A prefers for an U<u>C</u>a/u trinucleotide motif and A1+A1CF prefers A<u>C</u>u/a motif **(Fig. 2A)**, which is consistent with the reported motif preference for RNA editing by A3A and A1 **(*23, 29*)**. However, A3G did not show a clear motif preference here, possibly due to generally inefficient editing by A3G on a small number of edited sites (n=11) in this study **(Fig. 2A)**.

**Fig. 2.**
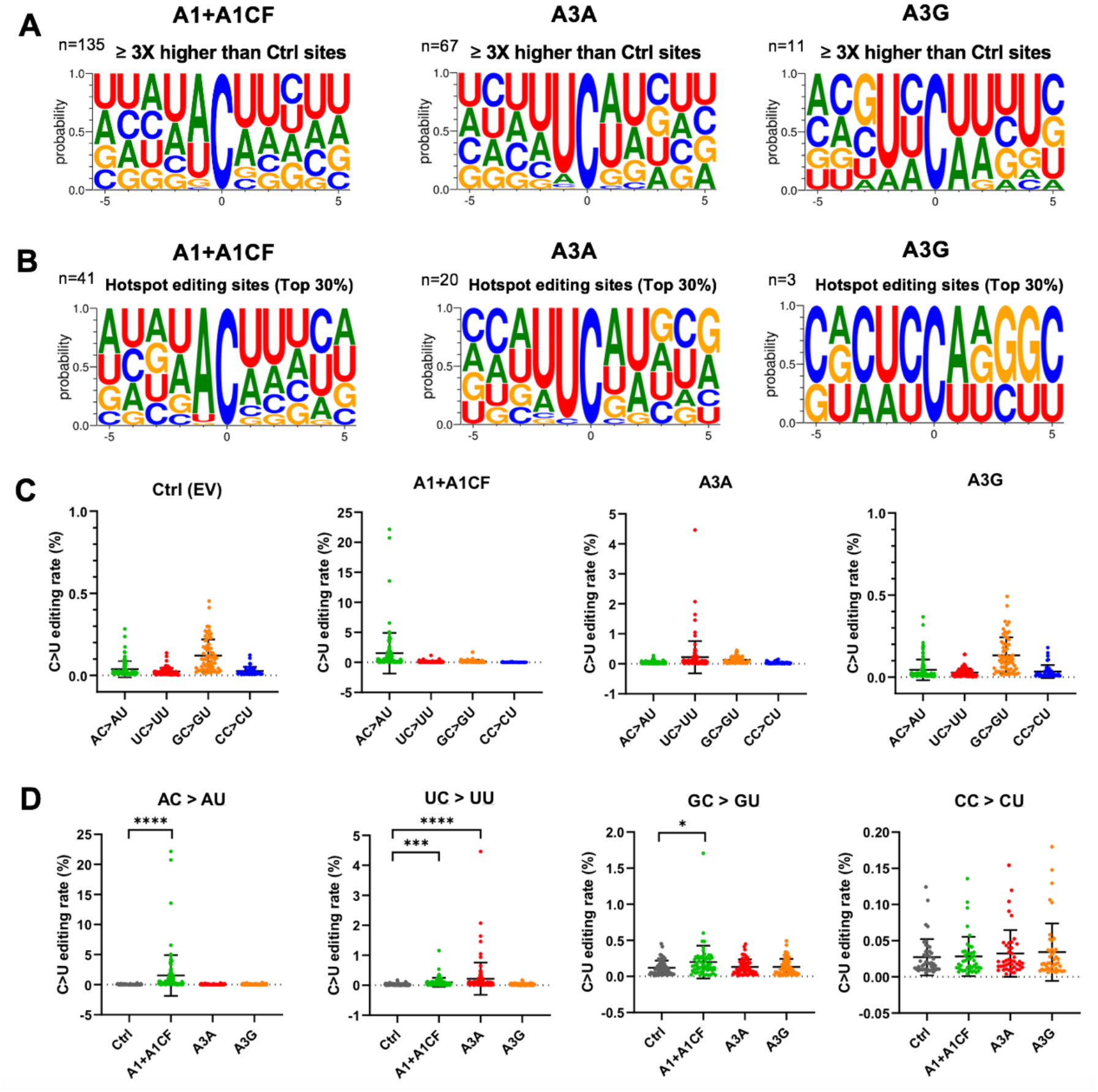
Local sequence context at the APOBEC-edited C sites on SARS-CoV-2 RNA. **(A)** Local sequences around the significantly edited target C sites (± 5 nucleotides from target C at position 0) by A1+A1CF, A3A, or A3G. The editing level of each C site was normalized to the Ctrl, and only sites with 3x or higher editing levels than the normalized value were defined as significant editing sites. **(B)** Analysis of local sequences around the top 30% edited C sites (or hotspot editing sites), showing predominantly A<u>C</u> motif for A1+A1CF, U<u>C</u> for A3A, and C<u>C</u> for A3G. **(C-D)** Comparison of the C-to-U editing rates (%) of different dinucleotide motifs by a particular APOBEC (panel-C) and the C-to-U editing rates (%) of a particular dinucleotide motif by the three APOBECs (panel-D). Each dot represents the C-to-U editing level obtained from the SSS results. In panel-D, statistical significance was calculated by unpaired two-tailed student’s t-test with *P*-values represented as: P > 0.05 = not significant; not indicated, * = P < 0.05, *** = P < 0.001, **** = P < 0.0001.

We also performed the same analysis of the sequence contexts with the top 30% editing efficiency by APOBECs (or hotspot editing sites), which is translated to 38 times higher than control for A1+A1CF, or 15 times higher than control for A3A, or 6 times higher than control for A3G) (**Fig.2B**). It distinctly shows that A3A strongly prefers the U<u>C</u> motif, whereas A1+A1CF has a strong bias toward the A<u>C</u> motif, and A3G prefers C<u>C</u> motifs on the viral RNA. These results suggest that the observed A<u>C</u>-to-A<u>U</u> mutations are most likely generated by A1 plus cofactor A1CF (or other A1 cofactors, such as RBM47 **(*30*)**); the U<u>C</u>-to-U<u>U</u> mutations by A3A, and C<u>C</u>-to-C<u>U</u> mutation by A3G in the sequence variation detected on the SARS-CoV-2 RNA segments (**Fig. 2C, D)**. Interestingly, among all the C-to-U variations of the SARS-CoV-2 sequences in our analysis, A<u>C</u>-to-A<u>U</u> (preferred by A1) and U<u>C</u>-to-U<u>U</u> (preferred by A3A) account for 38.23% and 31.83%, respectively, significantly higher than G<u>C</u>-to-G<u>U</u> and C<u>C</u>-to-C<u>U</u> that account for 15.44% and 14.50%, respectively **(Fig S2)**, indicating the significance of APOBEC editing on the viral mutation.

### Features of the efficiently APOBEC-edited RNA sites on SARS-CoV-2

Although each of the three APOBEC proteins showed a strong preference for specific dinucleotide sequence motifs (i.e. AC, UC, or CC sequence motifs) for editing on the viral RNA, the relative editing efficiency of these motif sites vary greatly, such as between 0.0041% to 22.15% for A1+A1CF, and between 0.0040% to 4.46% for A3A **(Table S2)**. Furthermore, many of these AC, UC, and CC motif sites on the viral RNA have no detectable editing by A1+A1CF, A3A, and A3G, respectively, suggesting that other RNA features beyond the dinucleotide sequence motifs, such as the secondary and tertiary structures, must play a role in the editing efficiency of a particular motif site.

The RNA editing by A1+A1CF was previously reported to require a so-called mooring sequence that has a general stem-loop structure around the target C and contains relatively high U/G/A content downstream of the target C (*31, 32*). However, this requirement for the mooring sequence and stem-loop structures are shown to be quite relaxed and still needs further characterization (*26, 33*). We analyzed the RNA features around the top 3 A<u>C</u> sites with the highest editing efficiency by A1+A1CF (**Fig. 3A**). The result showed that they could form a relatively stable stem-loop structure, with relatively high U/G/A contents downstream of the target C **(Fig. 3A)**. Among these top 3 editing sites, editing at C16054 is significantly higher than at C23170 (**Fig. 3A**), suggesting that, in addition to the possible involvement of long range RNA interactions, the editing efficiency also depends on the detailed local stem-loop structure and the position of the target C.

**Fig. 3.**
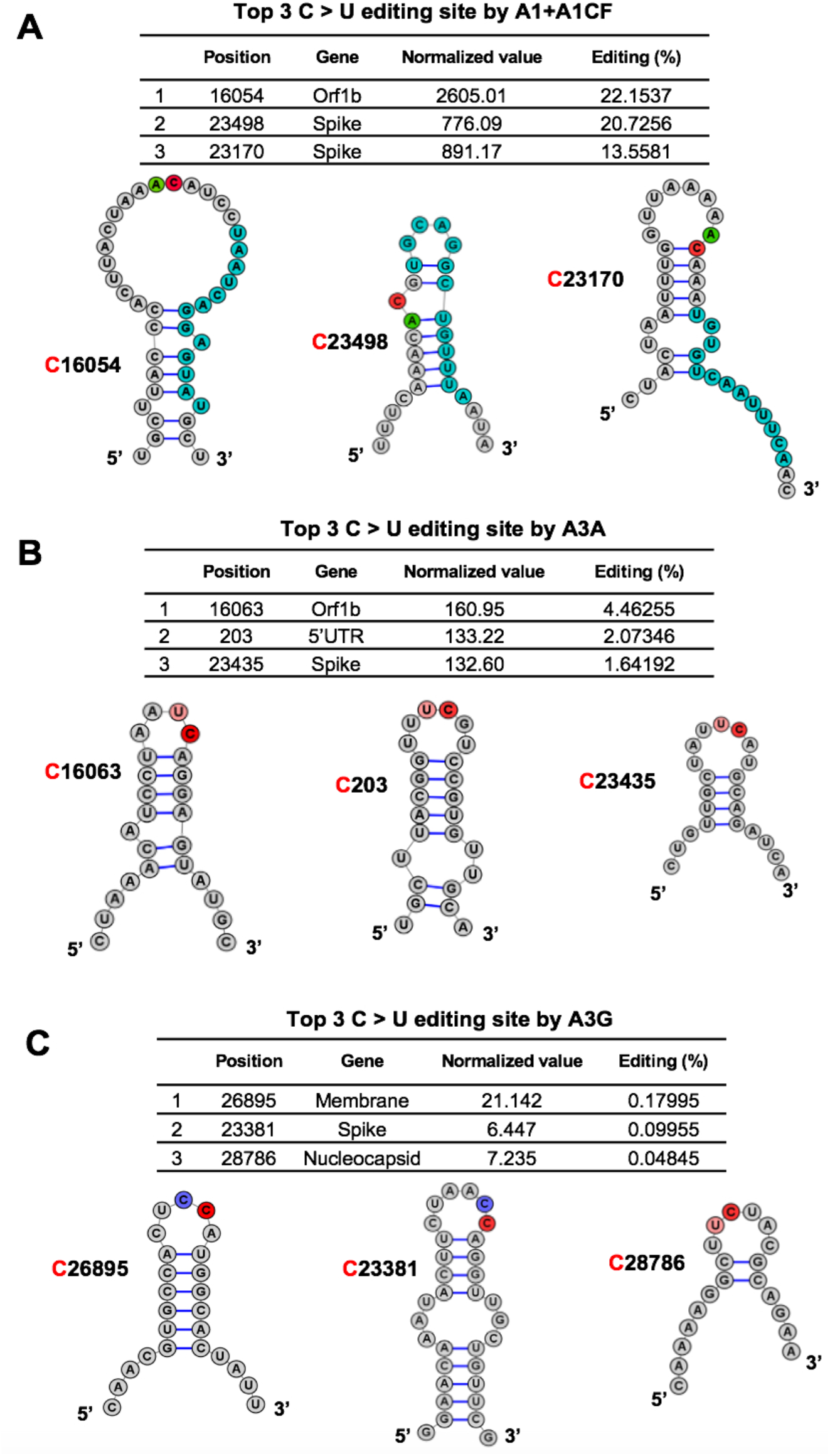
Overall features of the RNA around the most preferred APOBEC-edited sites on SARS-CoV-2. The predicted RNA secondary structures (*55*) of the sequences near the top 3 highest editing C sites by A1+A1CF **(A)**, A3A **(B)**, and A3G **(C)** (See related Table S2). The editing efficiency of each site is listed at the top of each panel. In the secondary structure, the target C sites are highlighted in red, and −1 positions of the target C sites are highlighted in green for A, pink for U, and blue for C, respectively. In panel-A, the proposed canonical mooring sequences for A1+A1CF (highlighted in sky blue) contain relatively high U/A/G contents downstream of the target C.

Sharma and colleagues recently discovered RNA editing activities by A3A and A3G in human transcripts through RNAseq using the common NGS without the safe sequencing SSS approach (*23, 24, 34*).More than half of target cellular RNA substrates have a stem-loop secondary structure, and the target C locates in the loop region. Interestingly, our top 3 highest A3A-mediated editing sites on SARS-CoV-2 RNA reported here all have the U<u>C</u> motifs in the loop of a predicted stem-loop secondary structure **(Fig. 3B)**. Again, the editing efficiency at C16063 (4.5%) is about 3-fold higher than the third efficiently edited site at C23453 (1.6%) **(Fig. 3B)**. Analysis of the top 3 sites edited by A3G also showed the CC (or UC) motifs in the single-stranded loop region of a predicted stem-loop secondary structure (**Fig. 3C**).

To rule out the possibility that the RNA C-to-U mutations result from DNA C-to-U deamination instead of the direct RNA editing by APOBECs, we performed a side-by-side sequencing of the DNA on the reporter vector and its corresponding RNA transcript containing the Orf1b region of SARS-CoV-2 (15,968-16,167 nt). The reporter DNA and the RNA transcript extracted from the cells expressing APOBECs were PCR amplified using the forward prime annealing to either the AAV-intron specific for the DNA or to the JUNC specific for the spliced RNA only. The PCR products from the DNA and mRNA were subjected to Sanger sequencing, and the C-to-U changes were analyzed. No DNA C-to-U mutation was detected, but specific RNA C-to-U changes on the mRNA transcript were present in a frequency consistent with our SSS results obtained in the presence of A1+A1CF (e.g., C16049, C16054, and C16092, etc.) or A3A (C16063) **(Fig S3)**. These data indicate that the C-to-U changes on the RNA are not caused by DNA mutation, but are the result of direct RNA editing mediated by A1+A1CF and A3A in our cell-based assay system.

### The potential effect of APOBEC-mediated RNA editing on SARS-CoV-2 variants

Because the current SARS-CoV-2 genome sequence databases are basically derived in the form of “one-consensus-sequence from one-patient”, they reflect the selected consensus viral sequences (including many of the mutations caused by APOBEC-mediated mutations) that survived the selection pressure for fitness. With the direct experimental evidence that APOBECs can target specific sites on SARS-CoV-2 for editing, we analyzed the publicly available SARS-CoV-2 genome sequence data (the Nextstrain datasets from Dec. 2019 to Jan. 22^nd^, 2022 downloaded from the GISIAD database, https://nextstrain.org/ncov/global)**(*35*)**, with hope to detect some obvious effects of APOBEC-mediated viral mutations on the current viral strain variants and fitness. The analysis revealed that the C-to-U is the predominant mutation for the entire genome, accounting for ~55.8% among all single nucleotide variants (SNVs) within the SARS-CoV-2 5’UTR-Orf1a region (142-341nt, the segment tested in our reporter 1 vector) **(Fig. 4A and Fig S4A)**. Of particular interest is the prominent mutation occurring at U**<u>C</u>**203, U**<u>C</u>**222, and U**<u>C</u>**241 in the 5’UTR region of these virus variants **(Fig. 4A)**, as these 3 sites all feature U**<u>C</u>** motifs and showed significant C-to-U editing by A3A in our assay results **(Fig. 4B, Fig S4B, and Table S2)**. These results suggest that A3A generated these mutations on the viral RNA genome, and the mutations can be maintained, likely because these three mutations generated by A3A editing are not determinantal to the virus. Two of these three mutations, U**<u>C</u>**203 and U**<u>C</u>**222, were detected in some of the SARS-CoV-2 sequences in late 2020 but are not persistently present in the main circulating strains, suggesting these two mutations may be neutral for the viral fitness. Surprisingly, the C-to-U mutation at U**<u>C</u>**241 occurred in early January 2020 and has rapidly become a signature of the dominant strains (including Delta and Omicron) that spread worldwide **(Fig. 4C and Fig S4C)**, strongly suggesting that this C-to-U mutation at U**<u>C</u>**241 may contribute to the better fitness for SARS-CoV-2. Although U**<u>C</u>**241to U mutation is within 5’UTR, the correlation of this mutation with the dominant new strains is reminiscent of that of the D614G mutation of the spike proteincoding region **(*5, 36*)**. Because 5’UTR has an important regulatory function for the replication of SARS-CoV-2 RNAs and for the expression of viral proteins **(*37, 38*),** the U**<u>C</u>**241 mutation may affect one or several aspects of these important functions of the 5’UTR in the viral infection steps relating to viral RNA replication, transcription, and translation.

**Fig. 4.**
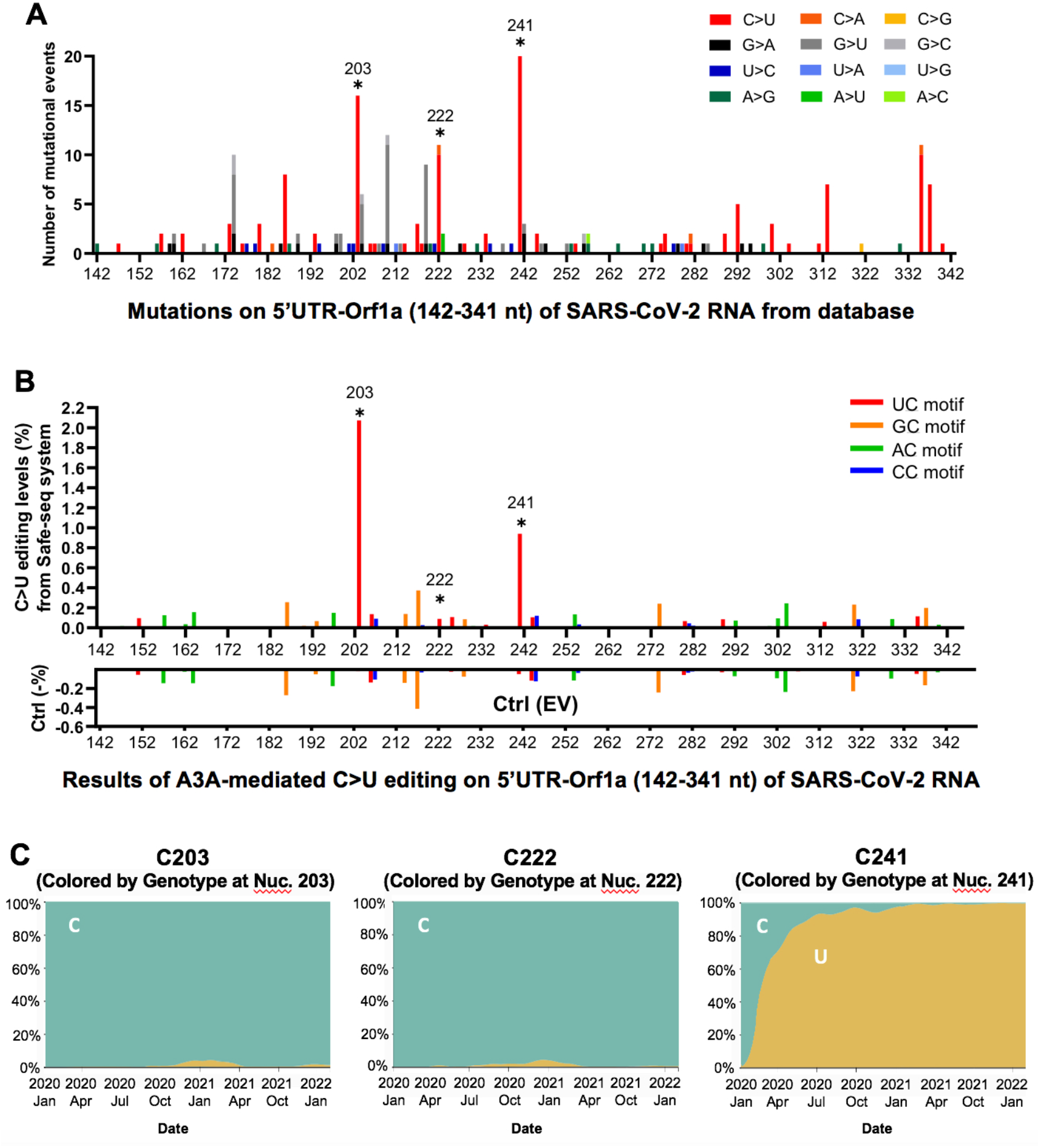
The potential effect of APOBEC-mediated editing on SARS-CoV-2 mutations and fitness. **(A)** The number of mutational events (all single nucleotide variants) on SARS-CoV-2 RNA segment 5’UTR-Orf1a (segment 1 in Fig. 1A) from the SARS-CoV-2 genome sequence data (the Nextstrain datasets from Dec. 2019 to Jan. 22nd, 2022 downloaded from the GISAID database, https://www.gisaid.org/hcov19-variants/ and https://nextstrain.org/ncov/global). The C203, C222, and C241 represent many of the C-to-U mutational events (asterisks) with the A3A-editing U<u>C</u> motif in the SARS-CoV-2 variants. **(B)** The A3A-mediated C-to-U editing rate on UC motif in the same 5’UTR-Orf1a region obtained from our cell-based editing system and the SSS analysis. The Ctrl (EV) editing levels (or background error rates) of the corresponding region are presented as negative values (%). The C203, C222, and C241 (asterisks) all showed significant editing by A3A. **(C)** The C-to-U mutation prevalence over time at C203, C222, and C241. The sequencing frequency is represented by C in blue and U in yellow (referred to the Nextstrain datasets: https://nextstrain.org/ncov/global). This analysis showed that SARS-CoV-2 started to acquire the C-to-U mutation at C241 in January 2020. By July 2020, 90% of the circulating viral variants carry this mutation at C241. By March 2021, almost all circulating viral variants have this mutation, suggesting the C241 to U mutation in the 5’UTR is beneficial to the viral fitness (see Fig. S9).

In all representative clades of SARS-CoV-2 emerged over the last two years since the initial outbreak, the C-to-U mutations have been much more pronounced than other types of single nucleotide variations **(Fig S5A)**. Even the very recent omicron variants, which began to spread from Nov. 2021 rapidly, continue to show noticeable C-to-U editing pattern, including some biologically significant mutations at the A1+A1CF preferred A<u>C</u> motifs **(Fig S5B)**. Notably, the A<u>C</u> to A<u>U</u> mutation at C23525 resulted an H655Y mutation on the spike protein**(Fig S5B)**, and the H655Y mutation was shown to alter cell entry pathways, i.e. the mutation is responsible for the preferential usage of endosomal pathway over cell surface entry pathways (*39*). This result suggests that further studies are necessary to have a comprehensive understanding of the potential effect of APOBEC-mediated C-to-U RNA editing on SARS-CoV-2 mutations and evolution.

### SARS-CoV-2 replication and progeny yield in cells overexpressing APOBECs

To examine if the three APOBEC proteins can affect SARS-CoV-2 replication and progeny yield in a well-controlled experimental setting, we used the human colon epithelial cell line Caco-2 that expresses ACE2 receptor and thus is a model cell line for SARS-CoV-2 infection and replication studies **(*40*).** Because Caco-2 cell lines have no detectible endogenous expression of A1, A3A, and A3G protein **(Fig S6)**, we first constructed Caco-2 stable cell lines expressing one of the three APOBEC genes. The externally inserted APOBECs are under tetracycline-controlled promoter so that their expression can be induced by doxycycline **(Fig. 5A and Fig S7A)**. The Caco-2-APOBEC stable cell lines were then infected by SARS-CoV-2, and the viral RNA replication and progeny yield were measured and compared with the control cell line without APOBEC expression. The viral RNA abundance as an indicator for RNA replication was measured using real-time quantitative PCR (qPCR) to detect the RNA levels using primers specific for amplifying three viral regions: the Nsp12 region, the S region, and the N region, covering the genomic and subgenomic regions. The viral progeny yield was assayed through plaque assay in Vero E6 cell line using the virions produced from the Caco-2-APOBEC stable cell lines at different time points post-infection **(Fig. 5A)**. Vero E6 cell line is highly sensitive to viral infection because of its defective innate immunity, allowing sensitive quantification of viral progeny produced from the Caco-2 cell lines.

**Fig. 5.**
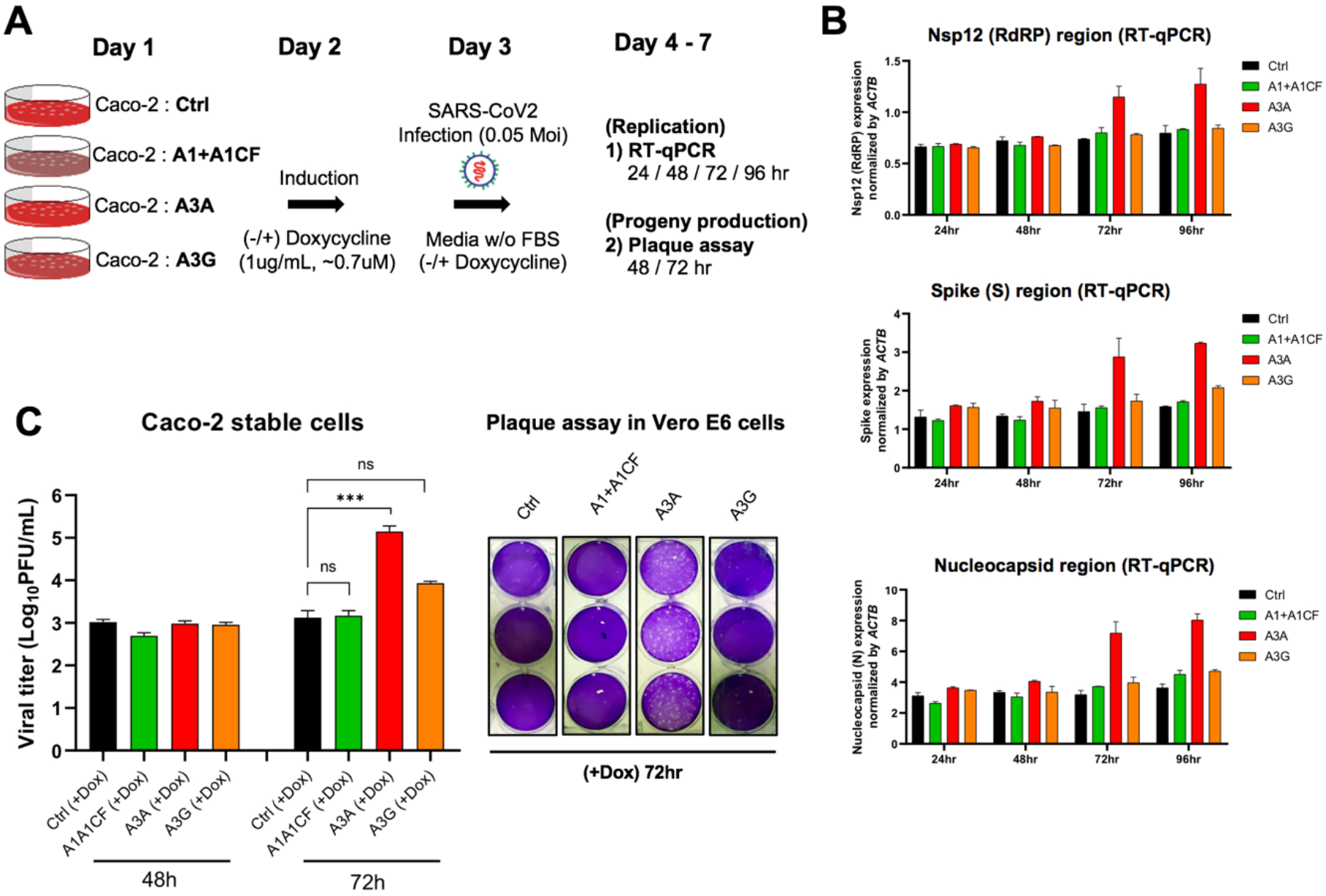
SARS-CoV-2 replication and virion production in cells expressing APOBECs. **(A)** Overview of experiments for SARS-CoV-2 replication and viral production in the presence of APOBECs. The Caco-2 stable cell lines were constructed to express A1+A1CF, A3A, or A3G under a tetracycline-controlled promoter. The Caco-2-APOBEC stable cell lines were then infected with SARS-CoV-2 (MOI = 0.05), and the viral RNA replication and progeny production were measured at different time points. **(B)** Effect of each APOBEC expression on SARS-CoV-2 viral RNA replication. Measurement of relative viral RNA abundance at different time points after viral infection of the Caco-2-APOBEC stable cell lines expressing A1+A1CF, A3A, or A3G. The viral RNA abundance was measured using real-time quantitative PCR (qPCR) to detect RNA levels by using specific primers to amplify three separate viral regions, the *Nsp12, S*, or *N* coding regions (see Methods in SI). **(C)** Effect of each APOBEC expression on SARS-CoV-2 progeny production. Infectious viral progeny yield harvested in the medium at 48 hrs and 72 hrs post-infection was determined by plaque assay (see Methods). Statistical significance was calculated by unpaired two-tailed student’s t-test with *P*-values represented as: P > 0.05 = not significant, *** = P < 0.001.

The replication assay results showed that, compared with the control Caco-2 cell line, no significant change of viral RNA level was detected by qPCR with the cell lines expressing A1+A1CF or A3G even up to 96 hours post-infection **(Fig. 5B)**. Unexpectedly, the abundance of viral RNAs has increased significantly in the cell lines expressing A3A at 72 hours and 96 hours post SARS-CoV-2 infection. These results suggest that, despite the general viral restriction function of APOBECs, presence of A3A expression appears to endow an advantage for the viral RNA replication.

Consistent with the increased viral RNA replication, A3A expression in the stable Caco-2 cell line also correlates with significantly higher viral progeny yield after 3 days of infection. While no difference in viral titer was observed at 48 hours from all cell lines with or without APOBEC expression, the virus titer from the cell line expressing A3A consistently showed an approximately 10 to 100 fold higher than the control and A1+A1CF and A3G expressing cell lines at 72 hours **(Fig. 5C)**.

Based on the intriguing results of A3A-related enhancement of SARS-CoV-2 replication and progeny virus yield, we further investigated further whether such effects are dependent on the deaminase activity of A3A. We constructed an A3A-knockout (ΔA3A) and an A3A catalytically inactive mutant-expressing (A3A-E72A) Caco-2 cell lines, and the SARS-CoV-2 replication and progeny yield were compared in the four different Caco-2 cell lines, i.e. original Caco-2 cell line (control), and Caco-2 with A3A-knockout (ΔA3A), and the stable cell lines expressing A3A WT or the inactive mutant A3A-E72A **(Fig S7B)**. Again, the abundance of viral RNA based on the qPCR results of the three viral regions (*Nsp12, S*, or *N* coding regions) all displayed significant increases in the cell lines expressing A3A-WT at 72 and 92 hours post SARS-CoV-2 infection but showed no significant difference in the control Coca-2 cells and the ΔA3A cells **(Fig S8A)**. Furthermore, the viral progeny yield harvested from the A3A-WT expressing cell line also was significantly higher than those from the control Caco-2 cells and the ΔA3A cells **(Fig S8B)**. Interestingly, A3A-E72A also showed slight enhancement of viral RNA replication **(Fig S8A)** and viral progeny yield **(Fig S8B)**. Taken together, this result suggests that the A3A deaminase activity plays a major role for promoting the viral replication and viral progeny production. This is consistent, among other things, with the observation that the U**<u>C</u>**241 to U**<u>U</u>**241 mutation, a site highly edited by A3A in our study, is within the viral packaging signal and near the viral replication regulation area at the 5’UTR **(Fig S9)**, which could explain why this mutation becomes prevalent in the widely circulating SARS-CoV-2 strains after January 2020 (**Fig. 4A-C)**.

## DISCUSSION

Several new viral strains have emerged in this ongoing COVID19 pandemic due to viral mutation. More viral strains are expected to evolve in the future due to continuous virus mutations. The added selection pressure from the clinical usage of vaccines and antiviral drugs could lead to new drug resistant and immune-escape viral strains. Prior bioinformatic analysis of the SARS-CoV-2 sequences suggested that some of the C-to-U mutations may be caused by APOBEC proteins instead of the random mutations caused by viral RNA polymerase during replication or ROS (*9, 10, 12, 25, 41, 42*). Here we described the first experimental evidence demonstrating APOBEC enzymes, A1+A1CF, A3A, and A3G, can target specific SARS-CoV-2 viral sequences for RNA editing, and the resulting mutations likely contribute to the viral replication and fitness.

We examined the C-to-U editing on selected SARS-CoV-2 RNA segments in HEK293T cells by A1+A1CF, A3A, and A3G in an APOBEC-RNA editing assay in a cell-based system (*26*) using the SSS safe sequencing approach (*27, 28*). The combination of the cell-based system and the SSS approach enable us to examine the individual APOBEC-mediated editing rate of specific viral RNA sequences, which could not be obtained from analyzing the currently deposited SARS-CoV-2 viral sequences that are derived as “one-consensus-sequence from one-patient”, thus, reflect only the final selected consensus viral sequences that survived the fitness selection.

The SSS approach also avoid the high-error rate associated with the conventional NGS sequencing used for obtaining the currently deposited SARS-CoV-2 viral sequences. Our results demonstrated that the three tested APOBECs show different editing rate at specific viral RNA sites, reaching as high as 22.2% for A1+A1CF, 4.5% for A3A, and 0.18% for A3G, respectively, on the selected viral RNA transcript populations **(Fig. 2)**. This editing efficiency at a specific site is a few orders of magnitude higher compared with the estimated ~10^−6^ – 10^−7^ random incorporation error rate for Coronaviruses replication (*43*). The data here demonstrate that A1 plus its cofactor A1CF can efficiently edit specific A<u>C</u> motif sties of the viral RNA **(Fig. 2-3)**. A high mutation rate of C in A<u>C</u> motif was also noted in SARS-CoV-2 and Rubella Virus by prior bioinformatic analysis (*3, 9, 25*). A3A showed a preferred dinucleotide U<u>C</u> motif, consistent with previous reports about the preferred motif for A3A-mediated editing of cellular RNA targets (*23, 44*).

However, the U<u>C</u> and A<u>C</u> motifs alone are not sufficient for efficient editing by A3A or A1+A1CF. Many of the U<u>C</u> and A<u>C</u> motifs in the SARS-CoV-2 RNA showed only the background level of C-to-U mutation **(Table S2)**, which suggest that additional RNA structural features around the UC/AC motifs should play an important role in dictating whether a U<u>C</u> or A<u>C</u> site can be efficiently edited **(Fig. 3)**. Prior reports show that some cellular RNA targets with stem-loop structures are favored by A3A and A1 editing (*23, 26, 32, 44*). However, the exact RNA secondary/tertiary structural features that can dictate the editing efficiency of a UC/AC site in the SARS-CoV-2 genomic RNA by A3A/A1 remains to be defined.

While the editing of SARS-CoV-2 RNA by A3A, A1+A1CF, and A3G has been demonstrated in our cell culture system, the analysis of the expression profiles reveals that A3A and A1+A1CF, but not A3G, are expressed in the human organs and cell types infected by SARS-CoV-2 **(Fig S10A, B)**. Such expression profiles make it possible for A3A and A1+A1CF to edit the viral RNA genome in the real world. Many human cell types expressing ACE2 in multiple organs can be infected by SARS-CoV-2, including (but not limited to) the lungs, heart, small intestine, and liver (*45, 46*). A3A expresses in lung epithelial cells, and, importantly, the A3A expression level is significantly stimulated by SARS-CoV-2 infection in patients **(*47–49*) (Fig S10A)**. A1 and its two known cofactors, A1CF and RBM47, are not expressed in the lungs but are expressed in the small intestine or liver **(*50*)** that can also be infected by SARS-CoV-2 **(Fig S10B)**.

A1 was not previously thought as a candidate which could edit the SARS-CoV-2 genome because of its target specificity and lack of expression in lungs, the primary target tissues of SARS-CoV-2 infection (*51*). In our experimental system, the editing efficiency of A1+A1CF on the A<u>C</u> motif is much higher than A3A on the U<u>C</u> motif **(Fig. 2C, D)**. Interestingly, analysis of SARS-CoV-2 variants from the database also showed higher A<u>C</u> motif mutations (38.3%) than U<u>C</u> motif mutations (31.2%) **(Fig S3)**, as also reported in the previous bioinformatic database analysis (*3, 9, 25*). These results indicate that many of the A<u>C</u>-to-A<u>U</u> mutations in the SARS-CoV-2 genome from patients can be driven by A1+A1CF mediated RNA-editing in the small intestine and liver with SARS-CoV-2 infection. Because the A<u>C</u>-to-A<u>U</u> mutations was not detected in A1 alone **(Fig S3)**, it indicates that the A<u>C</u>-to-A<u>U</u> editing also requires a cofactor A1CF. Given the closely overlapping RNA target and editing efficiency by another A1-cofactor RBM47 (*26, 30, 33, 50*), it’s likely that RBM47 may also enable A1 to perform A<u>C</u>-to-A<u>U</u> editing of SARS-CoV-2 RNA. Interestingly, both A1 cofactors A1CF and RBM47 were shown to physically interact with SARS-CoV-2 RNA in an interactome study (*52*), offering indirect evidence that these RNA-binding A1-cofactors could recruit A1 to target SARS-CoV-2 RNA for editing in the infected tissue.

Since APOBEC proteins play an important role in immune responses against DNA and RNA viral pathogens ((*17–20*) and references therein), we also investigated the possible effects of the three APOBECs on SARS-CoV-2 replication and progeny production in our experimental system **(Fig. 5A)**. Surprisingly, expression of WT A3A in the tested cells showed significant increases of the viral RNA replication and the viral progeny production after 72 hours postinfection **(Fig. 5B and 5C, Fig. S8)**, even though some increase of replication and progeny production was also observed in the inactive-A3A mutant expressing cell line **(Fig. S8)**. These pro-viral effect of A3A contrast to the known anti-viral effect of APOBECs (*17–20*) The data indicate that the deaminase activity of A3A to mutate the viral genome plays a critical role in the pro-viral effect even though a minor deamination-independent enhancement effect could not be ruled out.

While most APOBEC-mediated C-to-U mutations detected in our assay system may be detrimental to the virus and will be lost in the viral infection cycle, thus missing in the databank consensus viral sequence, some mutations beneficial to the virus’s fitness will be selected for in the new viral strains. If so, SARS-CoV-2 can then turn the tables on the APOBEC mutational defense system for its evolution, including (but not limited to) the improvement of viral RNA replication, protein expression, evasion of host immune responses, and receptor binding and cell entry.

Even though the mechanisms by which A3A-editing promotes SARS-CoV-2 replication and progeny production may be complex and require future investigation, one of the A3A-mediated mutations present in the current circulating strains could offer insights into how SARS-CoV-2 takes advantage of A3A-mediated mutations. Among many of the detected A3A edited sites on SARS-CoV-2 RNA in our study are three C-to-U mutations located in the 5’UTR region, U<u>C</u>203, U<u>C</u>222, U<u>C</u>241 (**Fig. 4A, B**). The SARS-CoV-2 from patients detected all of these three mutations at different stages since early 2020, but only major circulating viral strains since early 2020 acquired the mutation at U<u>C</u>241 (**Fig. 4C, Fig S4C)**, suggesting that C-to-U mutation at U**<u>C</u>**241 is selected for better viral fitness. This observation is surprising considering U<u>C</u>241 is in the non-coding 5’UTR region and, thus, not affecting the ORF coding of a protein, as in the case of the previously reported D614G mutation of the spike protein for viral fitness (*5, 36*). Therefore, this C241 to U mutation should not be related to a change of a viral protein function, such as cell surface receptor binding or polyprotein processing.

Mutations outside protein-coding ORFs of SARS-CoV-2 were previously designated as nonfunctional changes (*3*). However, the 5’UTR region of SARS-CoV-2 has important function in regulating protein expression and viral RNA replication and virion packaging (*37, 38, 53*). In fact, the U<u>C</u>241 is within the viral packaging signal sequence that has a stem-loop secondary structure which is in close proximity to the stem-loop structures involved in replication and leader transcription regulatory sequence (TRS-L) (*37*) **(Fig S9)**. Thus, this UC241 to U<u>U</u>241 mutation may impact the viral RNA packaging and virion production, and may even impact RNA replication, sub-genomic production, or/and the translation efficiency of the downstream proteins to endow the virus with better fitness.

There is also the possibility of host RNA or DNA editing by WT A3A that leads to increased viral RNA replication and viral progeny production. Given the known activity of A3A in mutating both cellular genomic DNA and RNA transcripts, the possibility of triggering certain cellular events by A3A editing/mutation activity that favors SARS-CoV-2 replication cannot be excluded and warrants future investigation.

In summary, we report here the experimental evidence demonstrating that certain sites of SARS-CoV-2 genomic RNA can be directly edited with high efficiency by A3A and A1+A1CF to cause C-to-U mutation. Critical factors dictating the RNA-editing efficiency by these two APOBECs include a dinucleotide motif U<u>C</u> for A3A or A<u>C</u> for A1 as well as certain RNA structural features around the target <u>C</u>. Even though some APOBECs, including A3A, are regarded as host antiviral factors, we show here that RNA editing of SARS-CoV-2 by A3A can promote viral replication/propagation. These results suggest that SARS-CoV-2 can take advantage of APOBEC-mediated mutation for their fitness and evolution. Unlike the random mutations caused by RNA replication or ROS, the presence of the finite number of the UC/AC motifs on the SARS-CoV-2 genomic RNA and the potentially correct prediction of the viral RNA structures suggest that it is possible to predict all the possible target C sites in both the coding and non-coding regions to be edited by these APOBECs. With the new selection pressure from use of vaccines and antiviral drugs and the continued large-scale circulation of SARS-CoV-2 variants among unvaccinated and vaccinated people, such prediction may be meaningful for anticipating potential new viral mutations and the emergence immune escape and drug-resistant strains.

## Acknowledgments

This work is supported in part by NIH AI150524 to X.S.C, and the startup fund to P.F. from Herman Ostrow School of Dentistry at the University of Southern California.

## Authors contributions

K. Kim, P. Calabrese and X.S. Chen designed the experiments; K. Kim, S. Wang, P. Calabrese, C. Qin, Y. Rao. performed the experiments and data analysis; K. Kim wrote the manuscript draft; P. Calabrese, S. Wang, P. Feng, commented the manuscript; and X.S. Chen revised the manuscript.

## Data and materials availability

All data is available in the manuscript or the supplementary materials. All materials used in the study are available to any researchers upon request.

## Supplementary Materials

Materials and Methods

Fig S1-S10

References (26–28, 35, 37, 40, 49, 56, 57)

Table S1

Table S2

Table S3

