## Supplemental Materials for "The Roles of APOBEC-mediated RNA Editing in SARS-CoV-2 Mutations, Replication and Fitness"

### Supplementary Materials

#### MATERIALS AND METHODS

##### The cell-based RNA editing system

The Cell-based RNA editing system is adapted from previously reported in reference (*26*). Briefly, reporter vectors containing DNA corresponding to the different RNA segments of SARS-CoV-2 (NC_045512.2) (see Fig. 1A, 1B) and the APOBEC (A1+A1CF, A3A, and A3G) editor vectors (see Fig. 1C) were constructed. A1+A1CF is constructed as one open reading frame (ORF) with a self-cleavage peptide T2A inserted between A1 and A1CF (A1-T2A-A1CF), which will produce individual A1 and A1CF proteins in a 1:1 ratio (*26, 54*). HEK293T cells were cultured in DMEM medium supplemented with 10% FBS, streptomycin (100 μg/mL), and penicillin (100U/mL) and maintained at 37 °C, 5% $CO_2$. One day before transfection, the cells (250 μL) were seeded at an approximate concentration of 250,000 cells/mL on an 8-well glass chamber (CellVis). The cells were then transfected with a mixture (25 μL) of an APOBEC editor vector (500 ng) and a SARS-CoV-2 reporter vector (50 ng) and 1.5 μL of X-tremeGENE 9 transfection reagent (Sigma) and incubated for 48 hrs. After harvesting the cells, RNA extraction with Trizol (Thermo Fisher) and DNA extraction with QuickExtract (EpiCentre) was performed, respectively, according to the manufacturer's recommended instructions. The editor and reporter vectors used in this study were listed in Table S3.

##### Sequencing library preparation

The extracted RNA was reverse transcribed with Accuscript High-Fidelity Reverse Transcriptase (Agilent) to produce the single-stranded cDNA using a specific primer annealing to the downstream sequence of SARS-CoV-2 reporter segments. The reaction was performed in a volume of 20μl containing 1μg of total RNA, 100 μM of reverse primer, 1X Accuscript buffer, 10 mM dNTP, 0.1M DTT, 8U RNase Inhibitor, and 1μl of Accuscript High-Fidelity Reverse Transcriptase (Agilent) for 1 hr at 42 °C. The cDNA was then amplified for 2 cycles by adding a forward primer annealing to the junction region (JUNC, Fig. 1B), where the AAV intron is spliced out. In this first 2-cycle PCR amplification, the forward and reverse primers were attached to barcodes consists of 15 randomized nucleotides as the Unique Identifier (UID), plus four tri-nucleotides designating four different experimental conditions: TGA for A1+A1CF; CAT for A3A; GTC for A3G; and ACG for Ctrl. Phusion® High-Fidelity DNA Polymerase (NEB) was used for this PCR reaction: 98 °C 5 min - (98 °C 30 sec, 71.4 °C 30 sec, 72 °C 1 min) x2 – 72 °C 5 min. This PCR product (330 bp) was then cleaned up using a spin column PCR cleanup kit (Thermo) to remove the free first-round barcode primers. The second-round PCR was performed for 30 cycles with Illumina flowcell adaptor primers using Phusion® High-Fidelity DNA Polymerase (NEB): 98 °C 5 min - (98 °C 30 sec, 72 °C 1 min) x30 – 72 °C 5 min. All 28 (4 editors x 7 different SARS-CoV-2 substrates) of the different pooled PCR products (399 bp) were combined in equal amounts for the final libraries. The final libraries were subjected to a full HiSeq Lane (PE150, 370M paired reads, Novogene). The primers for the sequencing library preparation were listed in Table S3.

##### Analysis of Safe-Sequencing-System

To distinguish a true mutation from random mutation during PCR and sequencing errors, we followed the approach as reported in (*27*). The details of our implementation of the method was described in (*28*). We wrote Python scripts to analyze the sequencing data. We only considered those sequencing reads such that (1) at least 85% of the bases matched the reference sequence, and (2) the quality scores for all the UID bases were 30 or greater (probability of a sequencing error < 0.001). We clustered reads with the same UID and barcode into UID families. We only considered those families with at least three reads with the same UID and barcode. At each nucleotide site, the mutation frequency is calculated by dividing a numerator by a denominator. The denominator is the number of UID families that, at this particular nucleotide site, have at least three reads with quality scores of at least 20 (probability of a sequencing error < 0.01; because of this quality restriction, the denominator may be different at different sites). The numerator is the number of UID families that, at this particular site, (1) have at least three reads with quality scores of at least 20, and (2) 95% of these reads have the same base, which is different than the reference. The probability that three out of three reads will all have the same sequencing error at a site is then $10^{-7}$ (=$(0.01^3)/(3^2)$).

**Caco-2 Stable cell line expressing APOBEC proteins**

We used lentiviral transfection to construct stable Caco-2 cell lines expressing A3A, A3G, and A1+A1CF to study the effect of APOBEC on SARS-CoV-2 replication because Caco-2 expresses the virus receptor ACE2 (***40***). Lentivirus was produced by lentiviral vector system pLVX-TetOne-Puro (Clon-tech) in HEK293T cells. The cells (about 2 x $10^6$ cells) were seeded in a 100 mm plate one day before transfection. The cells were then co-transfected with lentiviral packaging vectors, 1.0 µg of pdR8.91 (Gag-Pol-Tat- Rev, Addgene), 0.5 µg of pMD2.G (VSV-G, Addgene), and 1.7 µg of the pLVX-TetOne-Puro vector encoding the APOBEC proteins, using 20 µL of X-tremeGENE 9 transfection reagent (Sigma). Lentivirus-containing supernatant from infected HEK293T cells was collected after 70 hrs and filtered through a 0.45 µm PVDF filter (Millipore). Virions were precipitated with NaCl (0.3 M final) and PEG-6000 (8.5% final) at 4°C for 6 hrs and centrifugated at 4000 rpm at 4°C for 30 min. The pelleted virions were resuspended in 100 µL of MEM medium. Caco-2 cells (human colon epithelial cell line, ATCC) were cultured in MEM medium supplemented with 10% FBS, streptomycin (100µg/mL), and penicillin (100U/mL), and maintained at 37°C, 5% $CO_2$. The Caco-2 stable cell lines were generated by transducing with the lentivirus for 24 hrs and selected with 5 µg/ml of puromycin. The expression of A1+A1CF, A3A, or A3G was induced by adding 1µg/mL doxycycline for 24 - 96 hrs. Expression of these APOBEC proteins was verified by Western blot.

The **ΔA3A** Caco-2 cell line was created by puromycin selection after targeting N-terminus of genomic A3A exon-2 region with CRISPR-Cas9 methods (guide RNA sequence: UGGAAGCCAGCCCAGCAUCC) and inserting the SV40-promoter-Puromycin resistant gene (938 bp) through homology directed repair (HDR) system (left homology arm: 703 bp and right homology arm: 561 bp). A randomized guide RNA was used to generate a Caco-2 cell line as a negative control.

**SARS-CoV-2 virus replication and progeny production**

SARS-CoV-2 propagation, infection, and viral titration were performed as previously described (*56*). All SARS-CoV-2 related experiments were performed in the biosafety level 3 (BSL-3) facility (USC). For SARS-CoV-2 propagation, Vero E6-hACE2 cells were used. The cells were plated at 1.5 x $10^6$ cells in a T25 flask for 12 hr and infected with SARS-CoV-2 (isolate USA-WA1/2020) at MOI 0.005 in an FBS-free DMEM medium. Virus-containing supernatant was collected when virus-induced cytopathic effect (CPE) reached approximately 80%.

To assess the effect of APOBEC (A1+A1CF, A3A, and A3G) on SARS-CoV-2 RNA replication, the Caco-2-APOBEC stable cells (about 2 x $10^5$ cells) were plated in 12-well plates. After 15 hours, cells were treated or untreated with Doxycycline for 24 hours before infection. Before viral infection, the cells were washed with an FBS-free medium once. Viral infection was incubated on a rocker for 45 min at 37 °C. The cells were washed and incubated in a medium containing 10% FBS with or without Doxycycline. Total cellular RNA was extracted from the infected cells at 24, 48, 72, 96 hrs. Real-time quantitative PCR (qPCR) was used to quantify the viral RNA abundance level at the four different time points using viral RNA-specific primers to detect the Nsp12, S, and N regions. The qPCR of the internal actin RNA abundance level is used as a control by using actin-specific primers.

To assess the effect of APOBEC (A1+A1CF, A3A, and A3G) on SARS-CoV-2 viral progeny production, plaque assay was used on Vero E6-hACD2 cells that has defective innate immunity and is highly sensitive to viral infection, allowing sensitive quantification of viral progeny produced from the Caco-2 cell lines. Vero E6-hACE2 cells were seeded in 12-well plates. Once cell reached confluence, cells were infected with serially diluted SARS-CoV-2 virions collected from the infected Caco-2-APOBEC stable cells that express A1+A1CF, A3A, or A3G at 48 hrs and 72 hrs after viral infection. The medium was removed after infection, and overlay medium containing FBS-free 1 x DMEM and 1% low-melting-point agarose was added. At 48 and 72 h post-infection, cells were fixed with 4% paraformaldehyde (PFA) overnight and stained with 0.2% crystal violet. Plaques were counted on a lightbox.

**Quantitative real-time PCR**

Total RNA was extracted from the SARS-CoV-2 infected Caco-2 cells using Trizol (Thermo Fisher). The extracted RNA was then reverse transcribed with the reverse primers specific to Nsp12, S, and N coding regions of SARS-CoV-2, and b-Actin as an internal control, respectively, using the high-fidelity reverse transcriptase Protoscript II (NEB). The reaction was performed in a volume of 20 µl containing 1µg of total RNA, 100 µM reverse primer, 1X Protoscript II buffer, 10 mM dNTP, 0.1M DTT, 8U RNase Inhibitor (40U/µl), and 200U ProtoScript RT for 1 hr at 42 °C. Quantitative real-time PCR was then performed with SYBR Green (PowerUp™ SYBR™ Green Master Mix, Thermo Fisher Scientific) in a volume of 10 µl/well containing 1µl of reverse transcribed cDNA product from above, 0.25 µl of forward and reverse primers (10 µM), and 5 µl of PowerUp™ SYBR™ Green Master Mix (2X) using a CFX Connected Real-Time PCR machine (Bio-Rad). Primers used in this study were listed in Table S3. The indicated gene (*Nsp12*, *S*, *N*) expression levels were calculated by the 2-ΔΔCt method and normalized by b-Actin expression level.

**Western blot and antibodies**

For Western blot analysis, cells were lysed in 1x RIPA buffer (Sigma). Western blot analysis were performed from three independent transfections using FLAG-tagged APOBECs and HA-tagged A1CF. α-Tubulin: internal loading control. The lysates were then subjected to Western blot with anti-FLAG M2 mAb (F3165, Sigma, 1:3,000), anti-HA mAb (HA.C5, Abcam, 1:3,000), and anti- α-tubulin mAb from mouse (GT114, GeneTex, 1:5,000) as primary antibodies. Cy3-labelled goat-anti-mouse mAb (PA43009, GE Healthcare, 1:3,000) was subsequently used as a secondary antibody. Cy3 signals were detected and visualized using Typhoon RGB Biomolecular Imager (GE Healthcare).

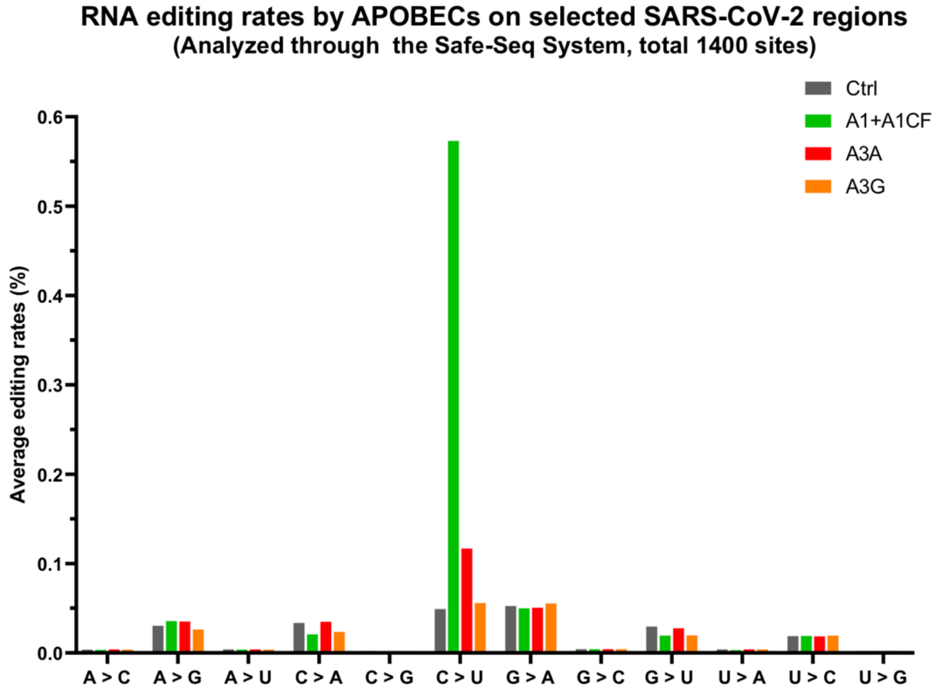


**Fig. S1.** The C to U RNA editing rates by APOBECs detected on the selected SARS-CoV-2 segments in our cell based assay system. Average rates (%) of all single nucleotide variations were analyzed through the Safe-Sequencing-System (SSS). See related Supplementary Table 1.

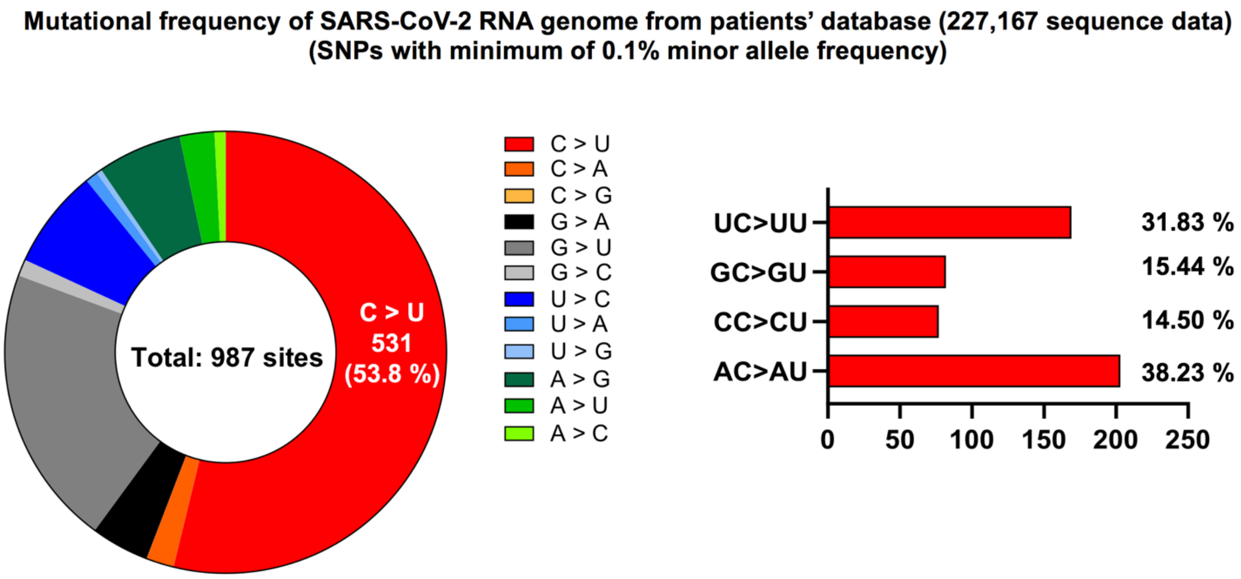


**Fig. S2. Single nucleotide variations (SNPs) of the SARS-CoV-2 genome sequences database derived from patients.** A total of 987 SNPs with minor allele frequencies > 0.1 % were counted from a total of 227,167 SARS-CoV-2 sequences on the UCSC genome browser (https://genome.ucsc.edu/covid19.html). The C-to-U mutation is the most common type with 53.8 % (left), of which the ratio according to the dinucleotide motifs including -1 position upstream of the mutated C are shown in the right chart.

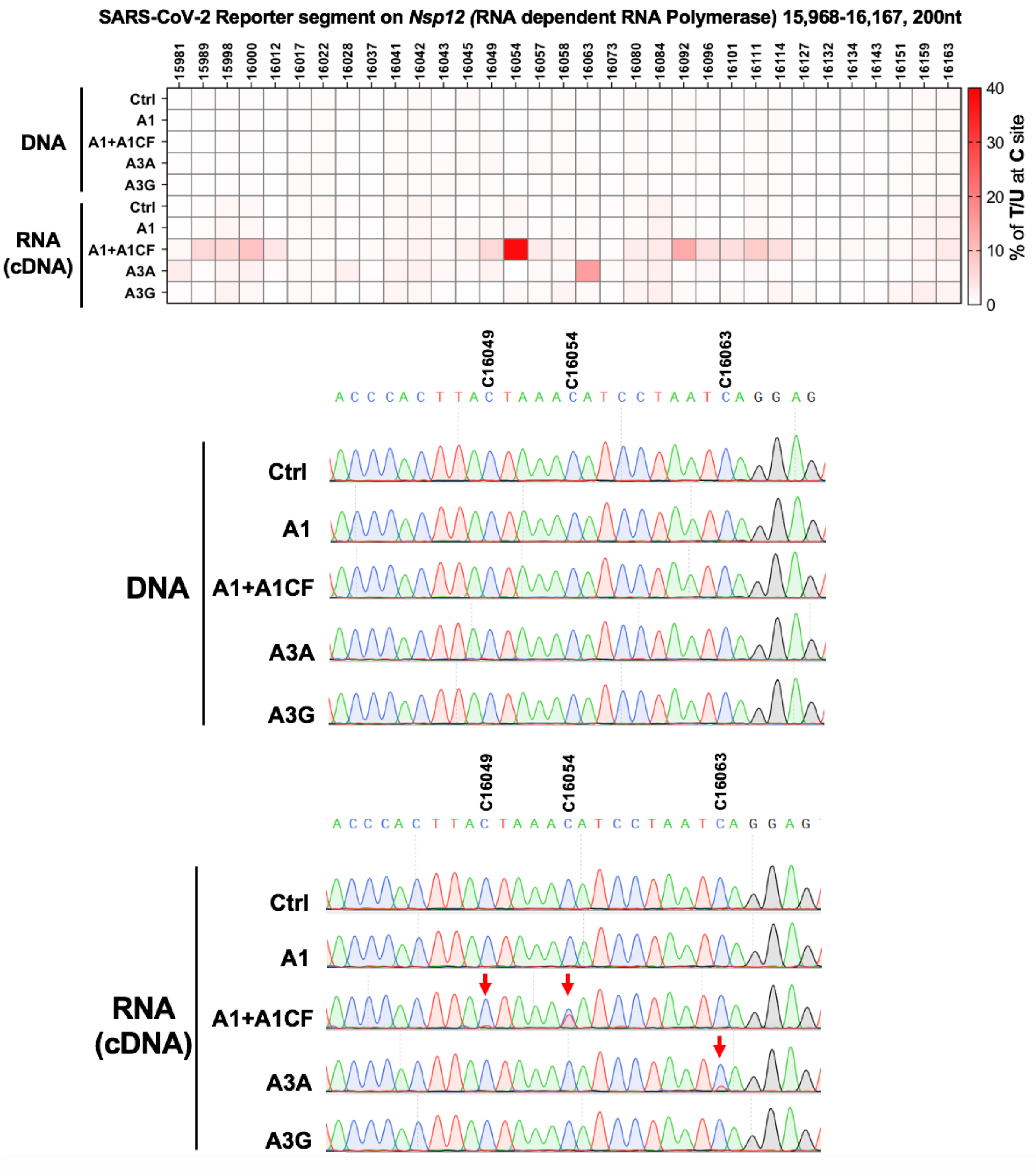


**Fig. S3. Verification of C-to-U mutation as a result of direct RNA editing on the transcript of a SARS-CoV-2 reporter segment (15968-16167 nt) instead of DNA deamination on the plasmid DNA by the three APOBECs A3A, A1 (+A1CF) and A3G.** The temperature-bar chart (top panel) shows the DNA and RNA C-to-T/U editing levels (%), which are based on the Sanger sequencing results of the DNA (middle panel) and the cDNA (RNA) (bottom panel). All C sites in this SARS-CoV-2 segment are marked with the virus nt sequence numbers on the top bar chart. three representative the RNA editing sites (C16049, C16054, and C16063) are indicated by red arrows.

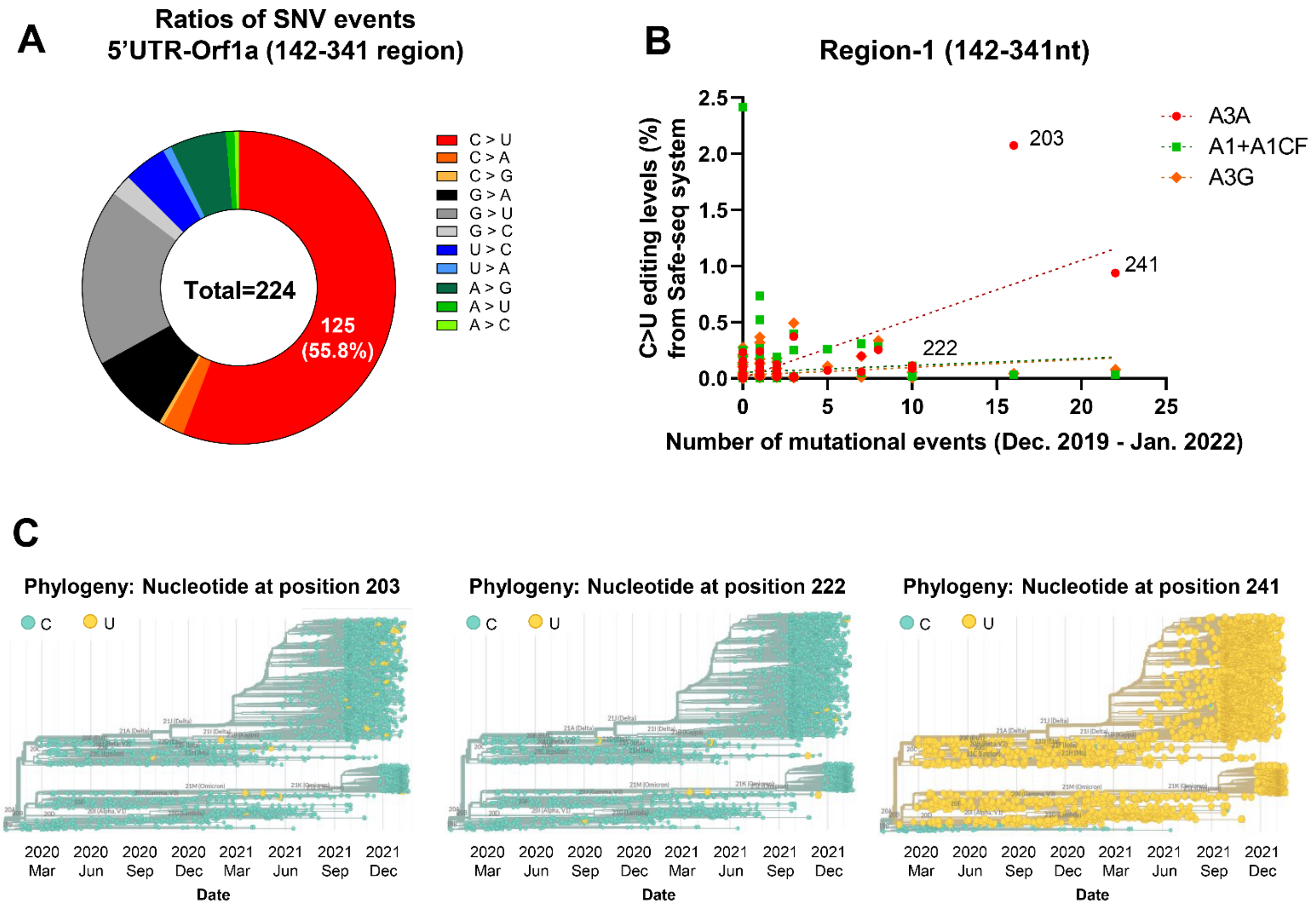


**Fig. S4.** Comparison of SARS-CoV-2 variants and the APOBEC-mediated RNA editing sites on the viral 5'UTR-Orf1a segment (142-341). **(A)** Ratios of all SNVs events on the 5'UTR-Orf1a segment from the sequence database (referred to the Nextstrain datasets (*35*)). : https://nextstrain.org/ncov/global) **(B)** Correlation of C-to-U RNA editing levels by the three APOBECs identified by our Safe-sequencing system (Y-axis) and the mutational events of SARS-CoV-2 from the sequence database between Dec. 2019 to Jan. 2022 (X-axis). Dotted lines indicate linear regressions with 95% confidence, and the case of A3A shows a positive correlation, and A1+A1CF shows a negative correlation. **(C)** Phylogenetic trees for C-to-U variant at C203, C222, and C241 (referred to the Nextstrain datasets (*35*)). : https://nextstrain.org/ncov/global). These phylogenetic trees correlate well with the C-to-U mutation prevalence over time at C203, C222, and C241, as shown in Fig. 4C.

**A**

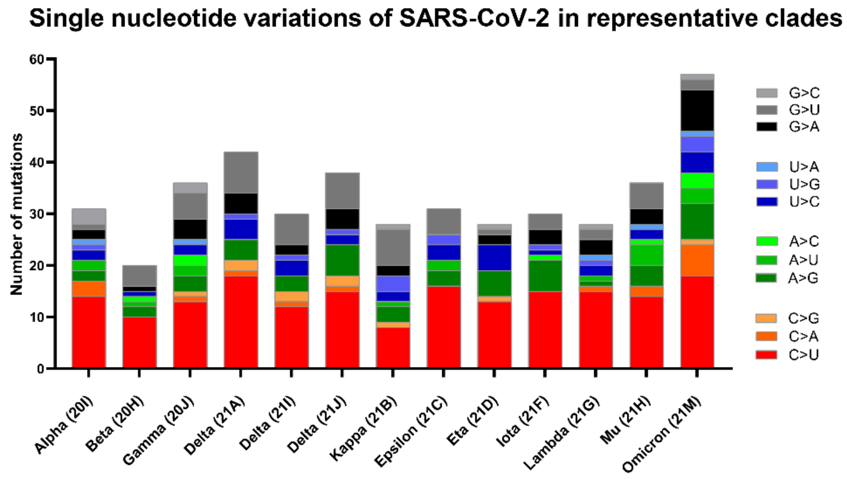


**B**

| Mutations | Gene | Frequency (2021 Jan - Dec) | Motif | Codon |
|---|---|---|---|---|
| C21U | 5'UTR | 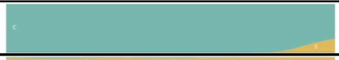 | C<u>C</u> | NA |
| C241U | 5'UTR | 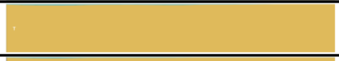 | U<u>C</u> | NA |
| C3037U | Orf1a (Nsp3) | 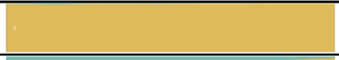 | U<u>C</u> | F924F |
| C10029U | Orf1a (Nsp4) | 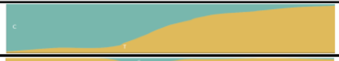 | A<u>C</u> | T3255I |
| C14408U | Orf1b (Nsp12) | 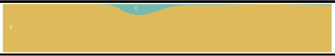 | C<u>C</u> | P214L |
| C15240U | Orf1b (Nsp12) | 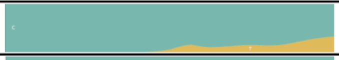 | A<u>C</u> | N491N |
| C21762U | Spike | 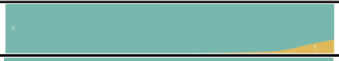 | G<u>C</u> | A67V |
| C21846U | Spike | 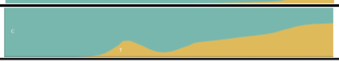 | A<u>C</u> | T95I |
| C22674U | Spike | 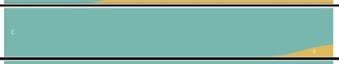 | U<u>C</u> | S371F |
| C22686U | Spike | 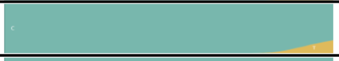 | U<u>C</u> | S375F |
| C23525U | Spike | 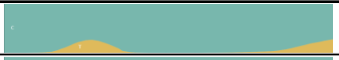 | A<u>C</u> | H655Y |
| C24503U | Spike | 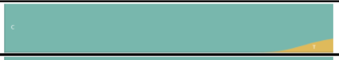 | C<u>C</u> | L981F |
| C25000U | Spike | 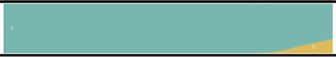 | A<u>C</u> | D1146D |
| C25584U | Orf3a | 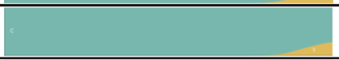 | C<u>C</u> | T64T |
| C26270U | Envelope | 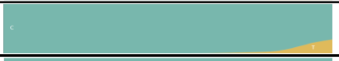 | A<u>C</u> | T9I |
| C27807U | Orf7/8 | 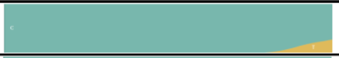 | U<u>C</u> | NA |
| C28311U | Nucleocapsid | 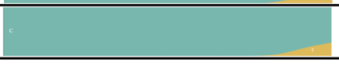 | C<u>C</u> | P13L |

**Fig. S5. Single nucleotide variations of SARS-CoV-2 in representative clades and characterization of C-to-U mutations in the *Omicron* variant (21M). (A)** Number of different single nucleotide variations (SNVs) in representative SARS-CoV-2 clades from *Alpha* (20I) to *Omicron* (21M). **(B)** Table listing the characterization of C-to-U mutations from the preferred editing motifs (U<u>C</u>, A<u>C</u>, and C<u>C</u>) by A3A, A1 (+A1CF), and A3G, respectively, in the representative *Omicron* variant (21M).

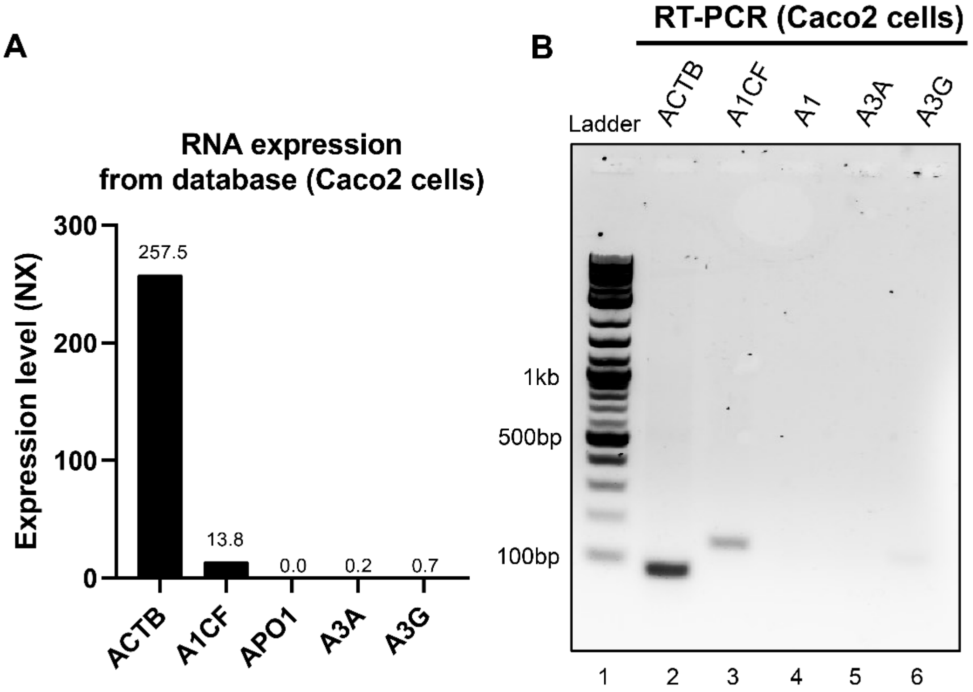


**Fig. S6. Examination of endogenous RNA expression of APOBEC editors in** the original **Caco-2 cells. (A)** Overall RNA expression levels of APOBECs (A1, A3A, and A3G) and A1CF in Caco2 cells from database. Each of gene expression values (NX) was retrieved from the human protein atlas (http://www.proteinatlas.org), which shows no or very low expression of these proteins. **(B)** RT-PCR analysis of the three APOBEC transcripts from the original Caco-2 cells. After RT of the total extracted mRNA, primers for detection of transcripts for β-actin (lane 2: 91 nt predicted), A1CF (lane 2: 139 nt predicted), A1 (lane 4: 160 nt predicted), A3A (lane 5: 143 nt predicted), and A3G (lane 6: 115 nt predicted) were used for amplification from the total cDNA. Lanes 2 was considered positive controls for genomic proteins of Caco-2 cells. This result is consistent with the analysis result from the proteinatlas database.

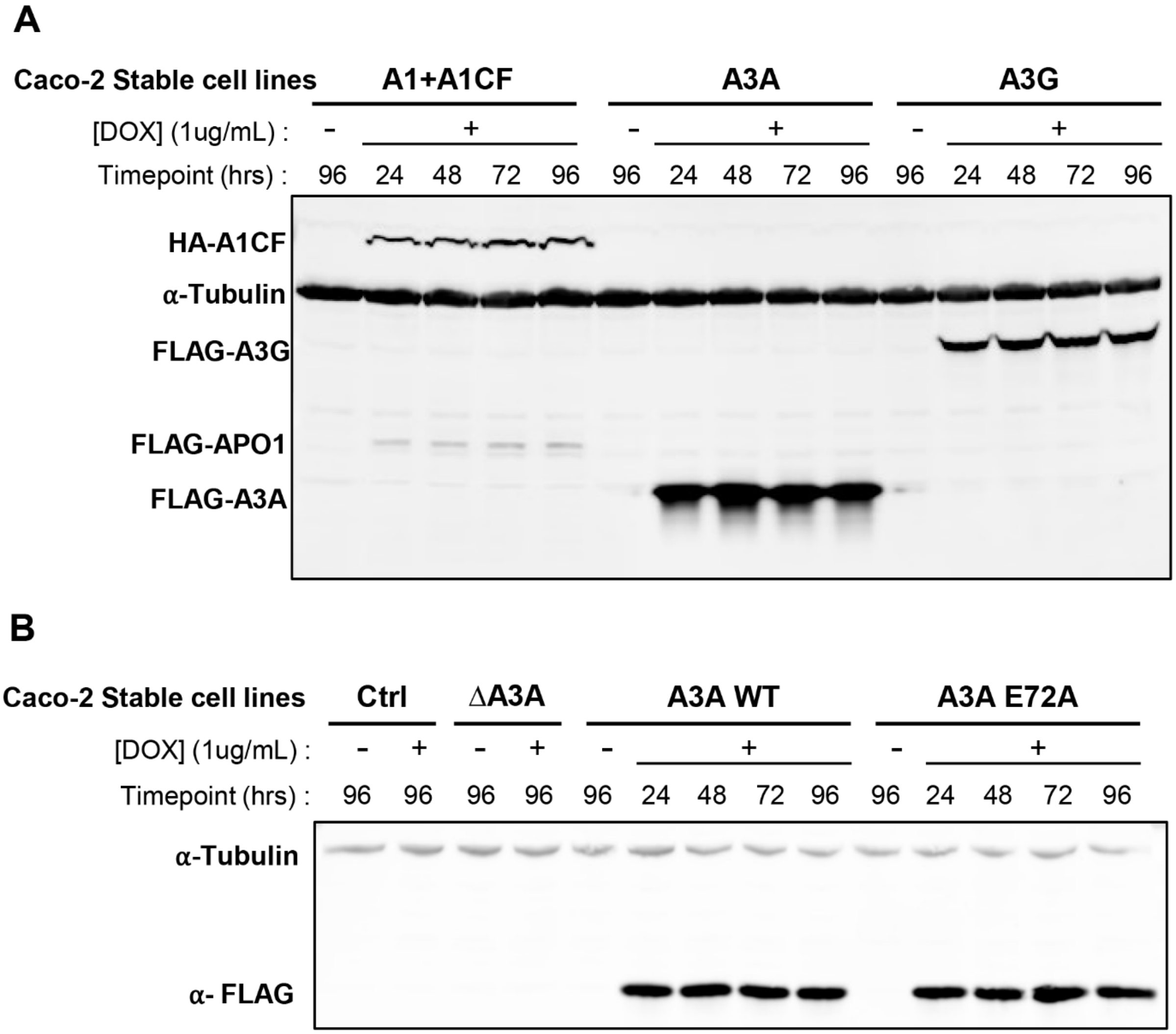


**Fig. S7. Western blot analysis of APOBEC protein expression from Caco-2-APOBEC stable cell lines.** **(A)** The expression level of the N-terminal FLAG-tagged A1 and HA-tagged A1CF (from A1-2A-A1CF construct), N-terminal FLAG-tagged A3A and A3G. **(B)** expression of N-terminal FLAG-tagged A3A WT and A3A E72A were detected under doxycycline treatment (1μg/mL) in different timepoints (24h, 48h, 72h, and 96h), whereas no A3A protein was detected in the A3A knockout caco-2 cell line and the control gRNA treated caco-2 cell line. α-Tubulin is the internal loading control.

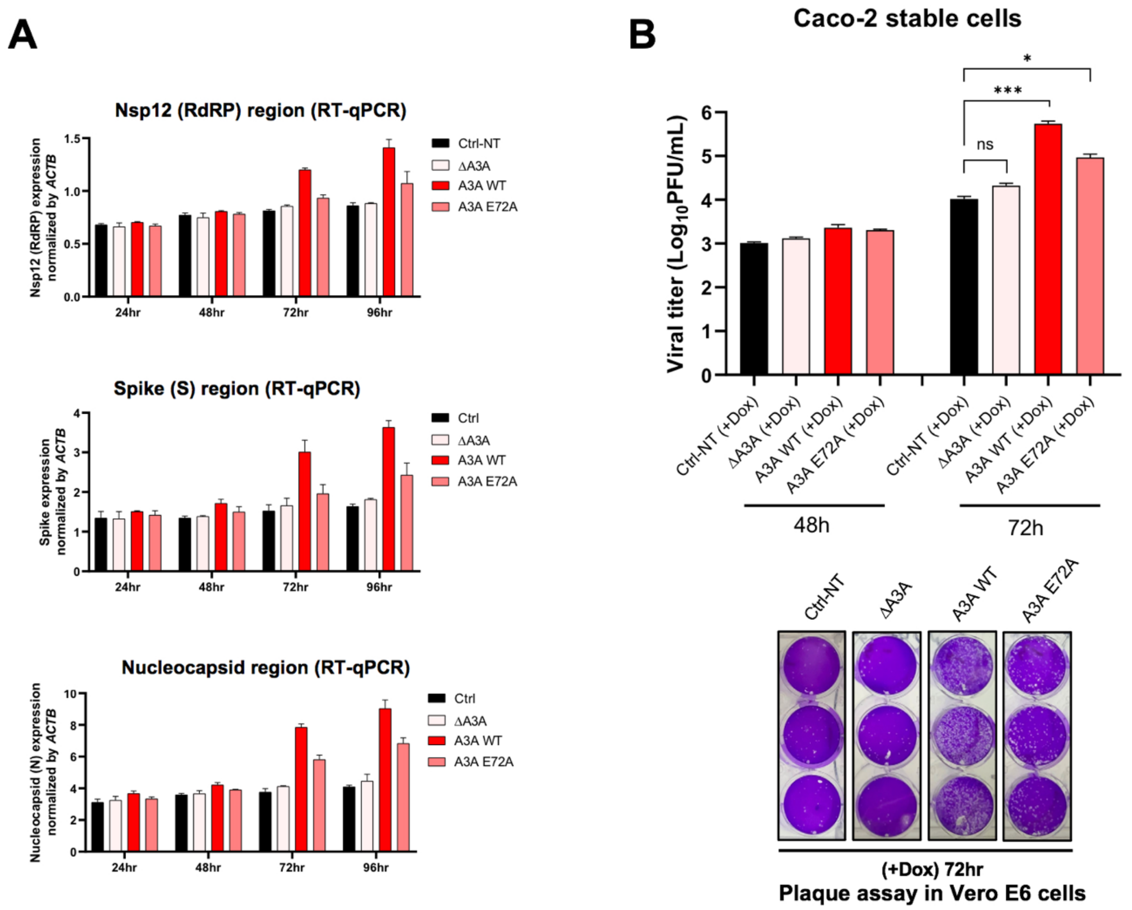


**Supplementary Figure 8. SARS-CoV-2 replication and progeny production in different Coca-2 cell lines** (Ctrl, ΔA3A, A3A WT, and A3A 72A)**. Ctrl:** randomized gRNA control Caco-2 cell line; **ΔA3A:** Stable Caco-2 cell line with A3A knockout by CRISPR; **A3A WT**: stable Caco-2 cell line expressing A3A wild-type protein; **A3A E72A:** stable Caco-2 cell line expressing catalytically inactive A3A mutant. **(A)** SARS-CoV-2 viral RNA replication in four different Caco-2 cell lines (Ctrl, ΔA3A, A3A WT, and A3A 72A). The viral RNA abundance was measured using real-time quantitative PCR (qPCR) to detect RNA levels by using specific primers to amplify three separate viral regions, the *Nsp12*, *S*, or *N* coding regions (see Methods). **(B)** SARS-CoV-2 progeny production in the four different Caco-2 cell lines (Ctrl, ΔA3A, A3A WT, and A3A 72A). Infectious viral progeny yield harvested in the medium at 48 hrs and 72 hrs post-infection was determined by plaque assay in Vero E6 cells (see Methods). Statistical significance was calculated by unpaired two-tailed student's t-test with *P*-values represented as: $P > 0.05$ = not significant, * = $0.01 < P < 0.05$, and *** = $P < 0.001$.

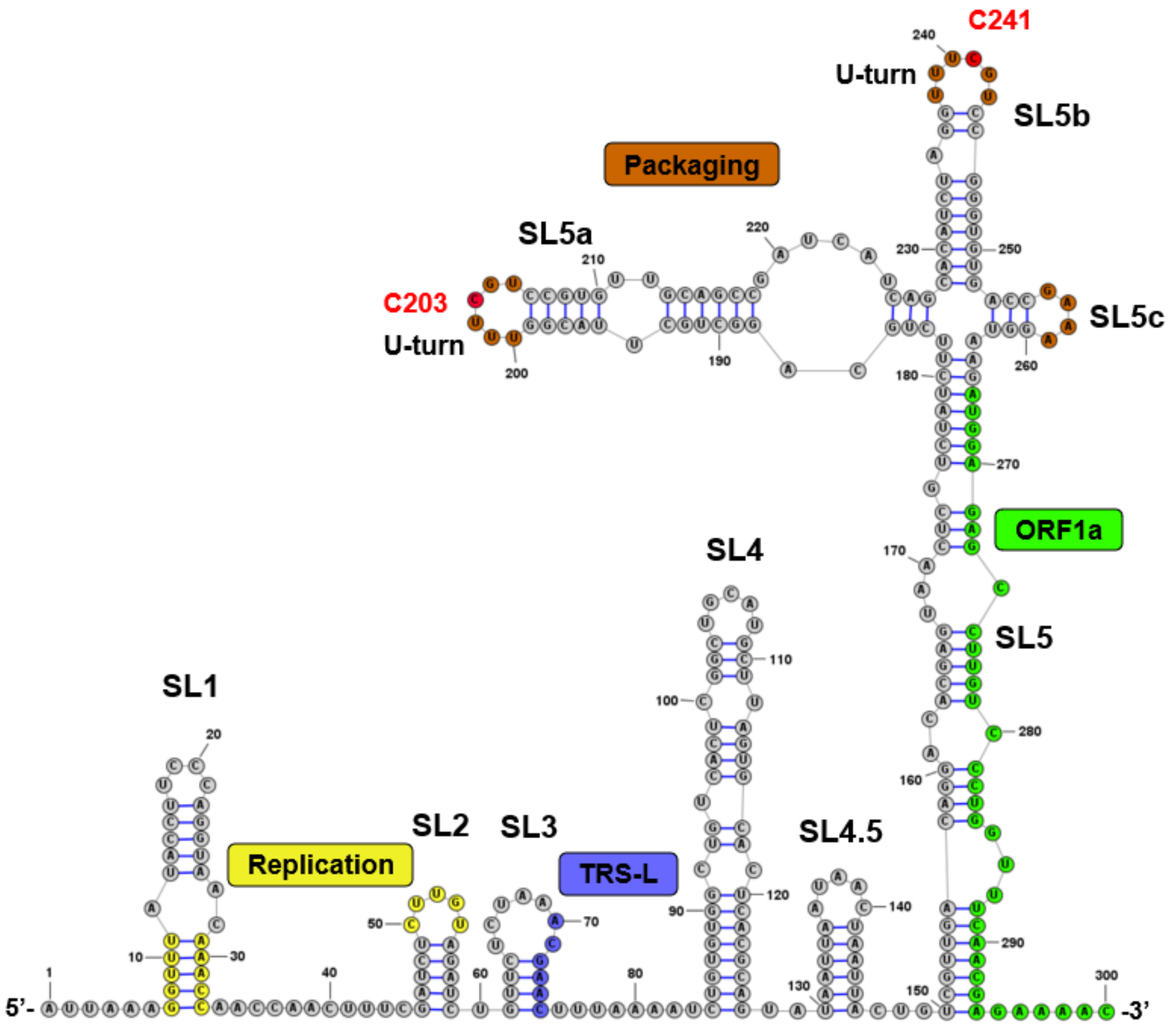


**Fig. S9. A predicted secondary structure for 5'UTR region of SARS-CoV-2 and its functional motifs.** The secondary structure model and functional motifs of SARS-CoV2 5'UTR were redrawn based on Miao *et. al.* (*37*). The packaging signals are highlighted in brown, replication-related motifs are highlighted in yellow, the leader transcription regulatory sequence (TRS-L) shown in blue, and ORF1a (from AUG) marked in green. The A3A editing target sites U<u>C</u>241 and U<u>C</u>203 (shown in red) are located on two separate loops within the packaging signal sequences that are spaciously close to the replication related motifs and TRS-L.

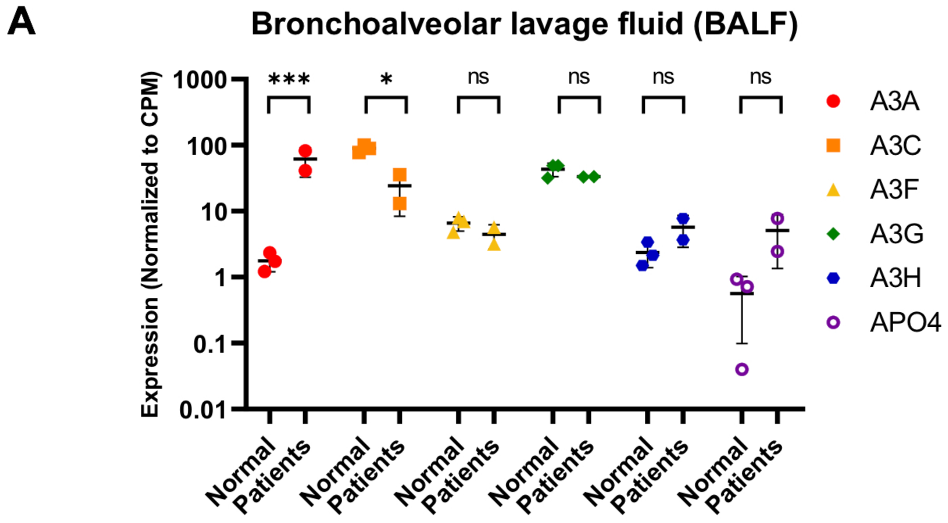


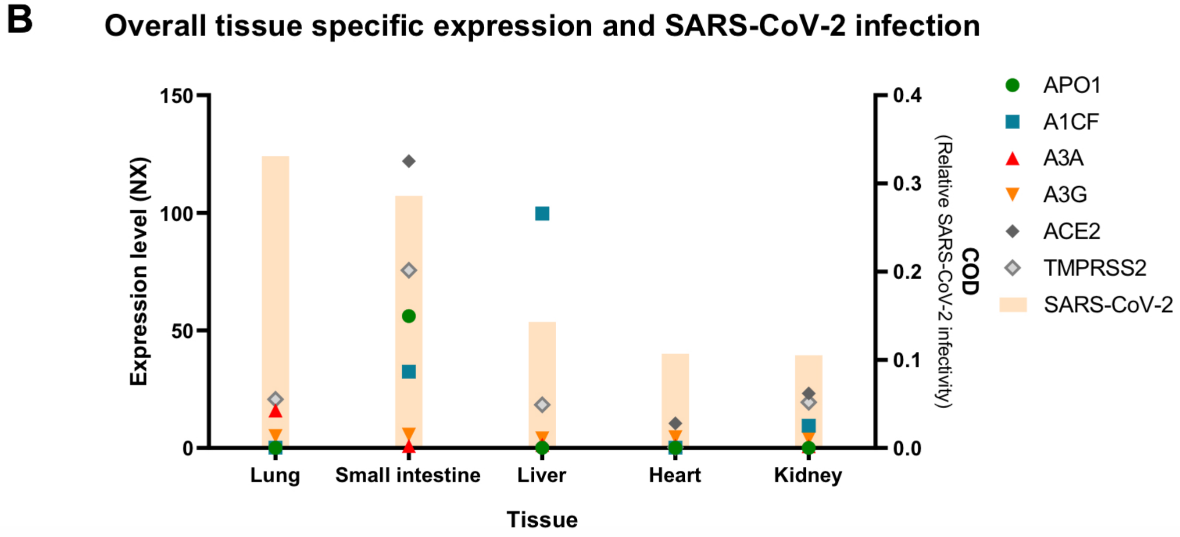


**Fig. S10. Relations between SARS-CoV-2 infection and APOBEC expression. (A)** Data analysis of the expression level of six APOBECs in healthy people and COVID-19 infected patients in Bronchoalveolar lavage fluid (BALF) samples (referred to the RNAseq data from reference (*49*)). **(B)** Overall gene expressions of the three APOBECs (A1, A3A, A3G) and A1CF in the tissues that can be infected by SARS-CoV-2. The commonness of viral detection (COD, relative SARS-CoV-2 infectivity) score for each tissue is indicated by yellow shaded boxes (referred to the COD score based on reference (*57*)). Each of gene expression values (NX) was retrieved from the human protein atlas (http://www.proteinatlas.org)
